# CASC15 dictates vascular smooth muscle cell growth fate and pathological vascular remodeling through post-transcription regulation of mitotic fidelity

**DOI:** 10.64898/2026.06.29.735301

**Authors:** Ibrahim A. Ahmed, Lakshanya Rajaganapathi, Santiago Rivero, Jianxin Wei, Sebastian Morales-Bermudez Espinel, Ariane Bruder, Angelina Kendi, Thiago Bruder-Nascimento, Cristina Espinosa-Diez, Delphine Gomez

**Author notes:** **<u>Corresponding author:</u> Delphine Gomez, PhD**, Associate Professor of Medicine, University of Pittsburgh Department of Medicine, Division of Cardiology, 200 Lothrop Street, Biomedical Science Tower, Room 1723, Pittsburgh, PA 15261.

## Abstract

Vascular smooth muscle cell (SMC) growth, whether hyperplastic or hypertrophic, is a central determinant of vascular remodeling in cardiovascular disease, yet the molecular regulators that direct SMC toward a specific growth fate remain poorly understood. Here, we identify the long non-coding RNA CASC15 as a critical regulator of SMC growth and vascular remodeling. CASC15 is enriched in the vasculature and SMC-rich tissues in humans and mice, and its locus harbors SNPs significantly associated with coronary artery disease and blood pressure. We identify a novel SMC-selective CASC15 isoform (S-CASC15) whose expression level determines SMC growth fate: elevated S-CASC15 promotes proliferation, while its loss drives hypertrophy, polyploidization, and binucleation. In vivo depletion of CASC15 limits vascular injury-induced neointima formation and atherosclerotic lesion expansion. Conversely, CASC15 overexpression exacerbates injury-induced neointimal hyperplasia. However, CASC15 KO mice exhibit spontaneous medial hypertrophy and vascular hypercontractility. Mechanistically, loss of S-CASC15 expression causes mitotic defects, followed by arrest in the G1 phase of hypertrophic and polyploid cells. We found that S-CASC15 pro-proliferative function is mediated through its interaction with RNA-binding proteins, including Nucleolin, and by regulating the stability of cell cycle checkpoint gene transcripts, thereby ensuring mitotic fidelity. Together, these findings establish CASC15 as a pivotal molecular switch governing the balance between hyperplastic and hypertrophic vascular remodeling and as a potential therapeutic target in cardiovascular disease.

## INTRODUCTION

Vascular remodeling is a central, evolutionarily conserved process occurring throughout the arterial vasculature, characterized by modifications in blood vessel caliber, structure, and cell composition. These morphological changes enable vascular adaptation but are also integral drivers of cardiovascular disease^1^. A major feature of both conductance and resistance artery remodeling is the involvement of vascular smooth muscle cells (SMCs), whose plasticity drives alterations in their phenotype and mass. SMC participation in vascular remodeling was originally described as a switch between a quiescent contractile state and a proliferative, non-contractile, synthetic state (referred to as phenotypic switching)^2,3^. However, it is now well established that this bistate model does not capture the complexity of SMC plasticity. SMC lineage tracing and single-cell transcriptomic studies have revealed the varied, context- and vascular territory-dependent phenotypic states SMCs exhibit in healthy and diseased vessels^4–7^. Similarly, increases in SMC mass within the vessel media or intima can be driven by distinct mechanisms: hyperplasia and hypertrophy. Hyperplasia, characterized by an increase in SMC mass due to cell proliferation, is a central feature of several major vascular diseases, including atherosclerosis, restenosis, and pulmonary hypertension^8–10^. Clonality mapping studies have demonstrated that SMC investment in atherosclerotic plaques and carotid artery neointima involves the oligoclonal expansion and hyperproliferation of a limited number of medial SMCs^11,12^. Conversely, seminal studies by Dr. Gary Owens’ group have provided compelling evidence that SMC hypertrophy primarily drives the increase in medial aortic SMC mass in hypertension^13,14^. Hypertrophy can lead to a 5-fold increase in SMC volume and has been associated with polyploidy. Interestingly, SMC hyperplasia and hypertrophy co-occur in hypertensive resistance arteries^15,16^. This suggests that, in a given pathological milieu, hyperplasia and hypertrophy are not mutually exclusive and may be triggered by common environmental cues. For example, Angiotensin-II (ANGII), a potent driver of hypertension, exerts context-dependent pro-proliferative and pro-hypertrophic effects on SMCs^17^. However, the molecular and cellular mechanisms that bias SMCs toward hyperplastic or hypertrophic remodeling remain partially understood.

SMC differentiation, contractility, and plasticity are tightly regulated by transcriptional and epigenetic mechanisms [reviewed in ^18^]. Many transcriptional programs play essential roles in the activation (e.g., SRF/Myocardin) or repression (e.g., KLF4) of SMC contractile genes, as well as in guiding SMC phenotypic transitions. Similarly, dynamic epigenetic mechanisms, including DNA methylation and histone modification landscapes, directly influence SMC phenotypic state through chromatin accessibility and transcription regulation. Moreover, histone modifications and histone variants have been implicated in the control of SMC lineage identity and plasticity^19,20^. In particular, our group recently established that the stable occupancy of H3K4me2 on the SMC gene repertoire is required for the maintenance of SMC lineage identity^19,21,22^. Gene-specific epigenome editing consisting of H3K4me2 demethylation on the Myocardin-regulated genes employing a Myocardin-LSD1 fusion protein revealed that H3K4me2 ablation was sufficient to induce impaired contractility, loss of lineage identity, and exacerbation of cellular plasticity in SMCs^19^. Importantly, this study also demonstrates that H3K4me2 is enriched at migration genes and primes their activation in response to environmental cues, ultimately enabling SMC migratory capacity and participation in vascular remodeling. Thus, the SMC-specific H3K4me2 signature serves as a priming and transcriptional memory mechanism allowing the dynamic activation, repression, and reactivation of genes involved in SMC contractile and migratory functions. H3K4me2 also mediates secondary SMC gene regulation by controlling microRNA (miRNA) expression^19,23^. The H3K4me2-editing mediated differential expression of miRNAs was predicted to predominantly contribute to SMC differential gene expression (60%)^23^. Among them, repression of miR145 contributed to impaired migration in H3K4me2-edited SMCs^19^. These studies highlight the cooperation and dependence between epigenetic mechanisms in regulating SMC phenotypic states. How maintenance or loss of H3K4me2-mediated SMC lineage identity influences the expression of other central non-coding RNA entities, including long non-coding RNAs, has not been evaluated.

LncRNAs act as signaling molecular complexes via their flexible and modular scaffolding properties. They facilitate diverse biological processes involving cell growth, splicing, morphology, cell differentiation, apoptosis and cell cycle^24,25^. Mechanistically, they can act epigenetically and transcriptionally through interaction with chromatin modifiers and post-transcriptionally by modulating miRNA bioavailability, and mRNA stability. High-throughput RNA sequencing (RNA-seq) revealed that lncRNA expression exhibits complex splicing regulation and a greater cell specificity than that of protein-coding genes, thereby serving as cell- and disease-state determinants^26–29^. Several evolutionarily conserved lncRNAs have emerged as regulators of SMC function and phenotypic states^30^. The lncRNA CARMN (cardiac mesoderm enhancer-associated non-coding RNA), first discovered during cardiac development^31^, is highly expressed in SMCs and promotes their differentiation by interacting with the transcription cofactor Myocardin (Myocd) to enhance transcription of the SMC contractile repertoire^32^. CARMN expression decreases in dedifferentiated SMCs upon pro-atherogenic stimuli and in human advanced atherosclerotic plaques. CARMN deletion or inhibition accelerates atherosclerosis development in mice^33^. SENCR (smooth muscle and endothelial cell-enriched migration/differentiation-associated non-coding RNA) also exhibits pro-differentiation effects, in part by acting as a sponge for the Myocardin-targeting microRNA, miR206^34,35^. In contrast, MALAT1 (metastasis-associated lung adenocarcinoma transcript 1), NEAT1 (nuclear paraspeckle assembly transcript 1), and SMILR (smooth muscle-induced lncRNA enhances replication) promote SMC dedifferentiation, proliferation, and atherogenic programs^36–39^. Adding to their growing translational and clinical relevance, Genome-wide association studies (GWAS) have identified cardiovascular disease risk variants within lncRNA loci^40,41^. Among the most robustly replicated findings, the Chr9p21 risk locus has emerged as the top GWAS signal for atherosclerotic cardiovascular disease since the first CAD GWAS in 2007, with risk SNPs within the gene body of the lncRNA ANRIL^42,43^. In the present study, we identify CASC15 as a novel, functionally relevant lncRNA that regulates SMC growth fate and whose locus harbors SNPs significantly associated with CVD^42,44–47^.

Cancer Susceptibility candidate 15 (CASC15) is a heavily spliced long non-coding RNA conserved across species, including humans, primates, and mice. CASC15 has been implicated in several types of cancers, with higher levels of expression generally associated with worse prognoses^48^. However, CASC15 effects appear to be highly context-dependent, as it has been reported to drive cell proliferation, cell migration, epithelial-to-mesenchymal transition, or cell apoptosis^49–53^. Several mechanisms by which CASC15 regulates the transcription and translation of proliferative and migratory signals have been identified, including interaction with chromatin remodelers (e.g., EZH2 and WDR5) and miRNA sponging^52,54–56^. Despite annotated CVD-associated SNPs in the *CASC15* locus, our understanding of CASC15’s role in the cardiovascular system and its relevance in CVD initiation and progression is remarkably limited. A study found that CASC15 expression promoted cardiomyocyte hypertrophy^57^. Supporting these conclusions, cardiac CASC15 expression was elevated after Transverse Aortic Constriction (TAC) surgery, and knockdown of CASC15 reduced ANG-II-mediated cardiomyocyte hypertrophy. Meanwhile, a recent study found that CASC15 promotes endothelial cell (EC) proliferation in pro-atherogenic conditions in vitro^58^. Yet, while most of the aforementioned studies establish the involvement of CASC15 in regulating multiple cellular functions across various tissues and pathological contexts, there is a general lack of characterization of the functional isoforms that drive these phenotypes. Furthermore, the potential relevance of CASC15 in vascular SMC has not been explored.

Herein, we identified CASC15 as a central regulator of SMC growth and participation in vascular remodeling. CASC15 was ranked as the most downregulated lncRNA in H3K4me2-edited SMCs compared with controls. CASC15 is enriched in the vasculature and SMC-rich tissues in humans and mice. We identified a novel VSMC-specific CASC15 isoform (S-CASC15), whose expression level biases the SMC growth fate towards hyperplasia or hypertrophy. While increased expression of this transcript variant enhanced SMC proliferation, its loss of expression drove cell hypertrophy, polyploidization, and binucleation in vitro and in vivo. The impact of CASC15 KO on vascular remodeling was several-fold: while reducing vascular injury-induced neointima and atherosclerotic plaque size due to lack of SMC investment, CASC15 KO caused baseline vascular and SMC hypertrophy, causing vascular hypervasoconstriction. S-CASC15 expression fluctuated during the cell cycle, peaking in the G2/M phase. S-CASC15 deficiency led to incomplete mitosis followed by arrest of hypertrophic and polyploid cells in G1. Mechanistically, S-CASC15 interacted with multiple RNA-processing regulators, including Nucleolin, and regulated the stability of G2/M and G1 gene transcripts. Together, this study established the central role of CASC15 in regulating SMC growth fate and biasing vascular remodeling toward hyperplastic rather than hypertrophic response through the regulation of cell cycle completion.

## RESULTS

### Loss of H3K4me2-mediated SMC lineage identity in SMCs alters lncRNA expression

We recently established the central role of the histone post-translational modification H3K4me2 in governing SMC lineage identity^19^. Locus-specific demethylation of H3K4me2 across the Myocardin-regulated gene repertoire led to profound alterations in the SMC transcriptomic profile. We evaluated the lncRNA expression profile in this bulk RNA-seq dataset, including H3K4me2-edited, PDGF-BB-treated, and control aortic rat SMCs (GEO: GSE179220) (**Supplementary Figure 1A**). Principal component analysis showed a clear separation among these three groups, with dispersion remarkably similar to that of total transcripts and miRNAs (**Supplementary Figure 1B**)^19,23^. H3K4me2 editing and PDGF-BB treatment were associated with the differential expression of multiple lncRNAs (**Supplementary Figure 1C-D**). We found 27 and 42 significantly differentially expressed lncRNAs in the PDGF-BB-treated and H3K4me2-edited SMCs, respectively, compared with contractile SMCs. 16 differentially expressed lncRNAs were similarly regulated in H3K4me2-edited and PDGF-BB-treated SMCs. Several lncRNAs implicated in the regulation of cardiovascular differentiation and function were identified in our screen. Fendrr (Fetal-lethal non-coding developmental regulatory RNA), essential for cardiac mesoderm differentiation in mouse^59^, was upregulated in the H3K4me2-edited SMC (**Supplementary Figure 1E**). Meanwhile, CARMN was strongly downregulated in both PDGF-BB- and H3K4me2-editing-induced SMC dedifferentiation. To investigate the putative functional relevance of other differentially regulated lncRNAs, candidates were prioritized based on their non-coding potential, evolutionary conservation between rodents and humans, and conservation of their genomic environment (**Figure 1A**). Among these candidates, *ABBR0727581.1,* ortholog of human *CASC15,* was the most downregulated lncRNA in H3K4me2-edited SMCs (**Supplementary Figure 1E**). *CASC15* is a gene adjacent to *SOX4* and is evolutionarily conserved in humans, primates, mice, and rats (**Figure 1B**). Importantly, *CASC15* proximal promoters as well as its first and second UTRs are highly conserved (**Supplementary Figure 2A, Supplementary Table 1)**. Together, we found that H3K4me2-mediated loss of lineage identity alters the lncRNA expression profile and identified CASC15 as a conserved lncRNA that is downregulated in epigenetically edited SMCs.

**Figure 1:**
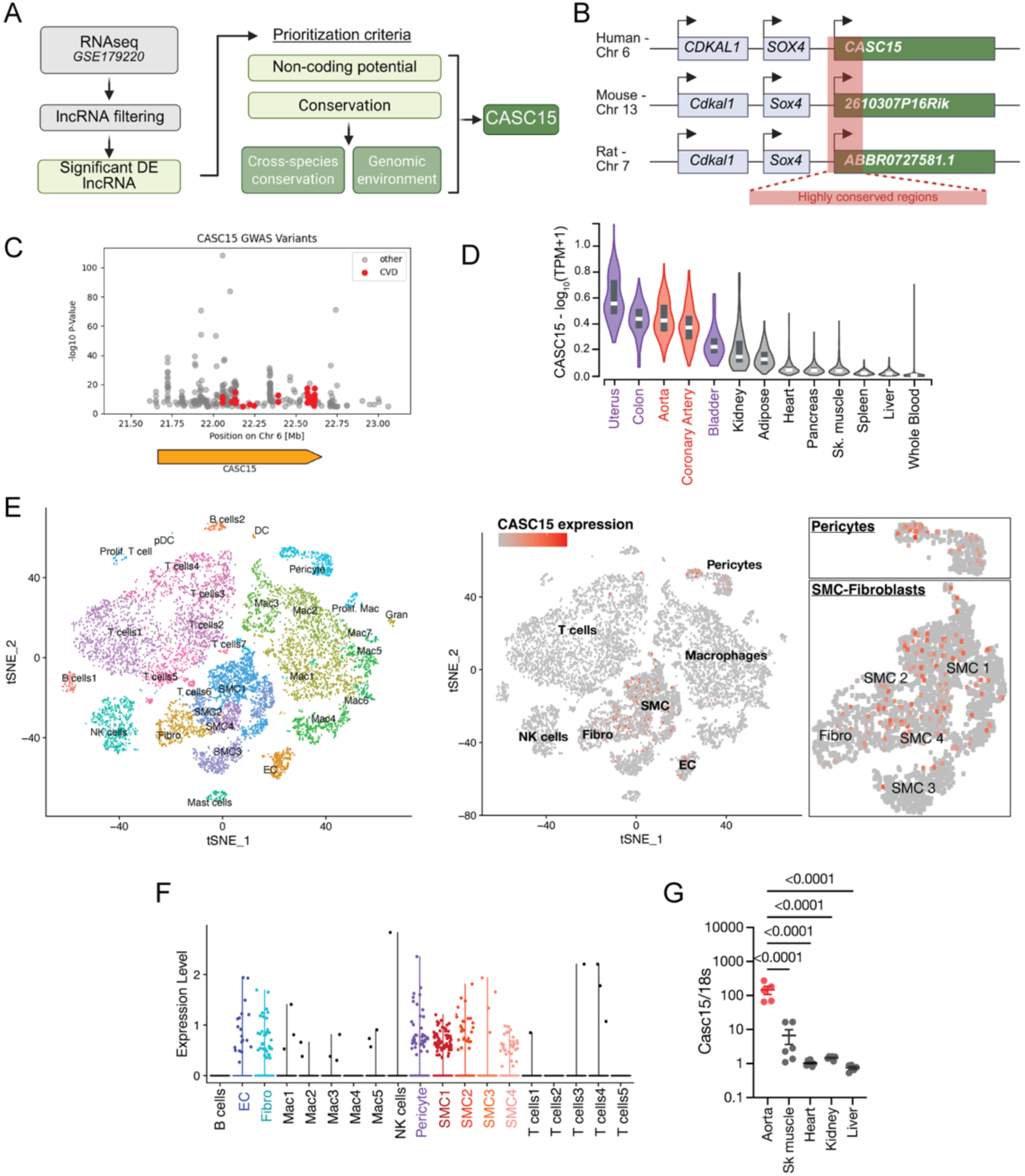
LncRNA CASC15 is preferentially expressed in SMCs. **A**. Schematics representing the analytical framework leading to the identification of CASC15 as a functionally relevant lncRNA candidate in SMC. **B**. Schematics the *CASC15* gene locus in human, mouse, and rat, highlighting evolutionarily conserved regions and conservation of the genomic environment across species. **C**. Regional Manhattan plot mapping SNPs with p<10^-8^ in the CASC15 locus. SNPs associated with cardiovascular traits and diseases are highlighted in red. **D**. CASC15 expression in human tissue (source: GTEx). Red: vasculature; purple: SMC-rich tissue. **E.** t-Distributed Stochastic Neighbor Embedding (t-SNE) and clustering based on single-cell RNAseq in human aorta (left). t-SNE plot for CASC15 expression (right). **F.** CASC15 expression levels in cell clusters identified in E. **G**. CASC15 expression in mouse tissues. One-Way ANOVA.

### LncRNA CASC15 expression is enriched in the vasculature and SMC-rich tissues

CASC15 function in the vasculature, and vascular SMCs in particular, has not been investigated. However, SNPs within and near *CASC15* have shown significant association with cardiovascular traits and diseases^42,44,46,60^. Integration of GWAS studies demonstrates the presence of SNP clusters (p<10^-8^) in *CASC15* associated with blood pressure and coronary artery disease (**Figure 1C**, **Supplementary Table 2-3**). Secondary analyses of publicly available datasets were performed to characterize CASC15 expression. CASC15 was found to be enriched in the vasculature and SMC-rich visceral and reproductive organs (i.e. colon, bladder, uterus) (**Figure 1D**). The CASC15 expression across tissues was comparable to the SMC-enriched lncRNAs SENCR and MYOSLID, but was not restricted to vascular SMC, in contrast with SMILR (**Supplementary Figure 2B**). Next, we evaluated CASC15 expression across vascular cell types by analyzing a published single-cell RNA-seq dataset from human ascending aortas (GSE189795). CASC15 expression was clearly enriched in SMC clusters (**Figure 1E-F**). CASC15 expression was also detected in pericytes, fibroblasts, and endothelial cells, but was lacking in immune cells. A similar pattern of CASC15 expression was found in human coronary arteries (GEO: GSE131780) and mouse tissues (**Figure 1G**, **Supplementary Figure 3A**)^5^. Interestingly, elevated CASC15 expression was detected in fibromyocytes, presumably derived from resident SMCs, in human atherosclerotic lesions. Concordant with these observations, single-nucleus ATACseq revealed greater chromatin accessibility on the *CASC15* gene in SMC and fibroblast clusters in coronary arteries from patients with CAD (GEO: GSE175621) (**Supplementary Figure 3B**)^61^. These data provide strong evidence for basal expression of CASC15 in human vascular SMCs and SMC-rich tissues, as well as enhanced expression in SMCs populating atherosclerotic plaques, further motivating the investigation of CASC15 function in SMCs.

### SMCs express a novel lineage-selective CASC15 isoform

CASC15 transcripts are heavily spliced. 267 annotated transcripts originating from mouse *2610307P16Rik* (chr 13:28,460,034-28,885,422) encoding CASC15 has been annotated with to date. Meanwhile, over 300 CASC15 isoforms have been characterized in humans. Using Alternative Splice Site Predictor (ASSP)^62^, we identify putative splice sites in *2610307P16Rik* transcripts (**Supplementary Table 4**). Based on these putative splicing sites and systematic untranslated exon amplification, we performed 5’-3’ end-point PCR and identified two novel CASC15 isoforms expressed in SMC, which have not been previously annotated (**Figure 2A**, **Supplementary Figure 4**). An 896bp short isoform (S-CASC15) contained untranslated exons 2, 9, 25, 29, and 33. A 1227bp long isoform (L-CASC15) was composed of a truncated untranslated exon 2, followed by untranslated exons 9, 25, 29, 33, 54, 56, and 57 (**Figure 2A-B**). Structural modeling with the Vienna Suite and AlphaFold3 revealed differences between the structures of these two isoforms, suggesting they may have distinct functions (**Figure 2C**, **Supplementary Figure 5A**). Performing isoform-specific qPCR, we found that S-CASC15 was selectively expressed in SMC, with minimal to no expression in endothelial cells, cardiomyocytes, and valve interstitial cells (**Figure 2D, Supplementary Figure 5B**). In contrast, L-CASC15 was more broadly expressed, with high expression in cardiomyocytes. Finally, we found that S-CASC15 was dominant in SMC compared with L-CASC15 (**Figure 2E**). CASC15 isoform expression is also tissue-selective in humans (**Supplementary Figure 6**). High sequence homology and conservation were observed between our newly identified mouse S-CASC15 and the human CASC15 untranslated exon 23-28 junction, which is present in multiple annotated isoforms (**Figure 2F-G**). In human aortic SMC (HASMC) RNA extracts, the E23-28 junction was amplified, and CASC15-208 was identified as the dominant mouse S-CASC15 equivalent (**Figure 2 H-I**). Given their conservation, SMC specificity, and expression profile, the functions of mouse S-CASC15 and human CASC15-208 were further investigated.

**Figure 2:**
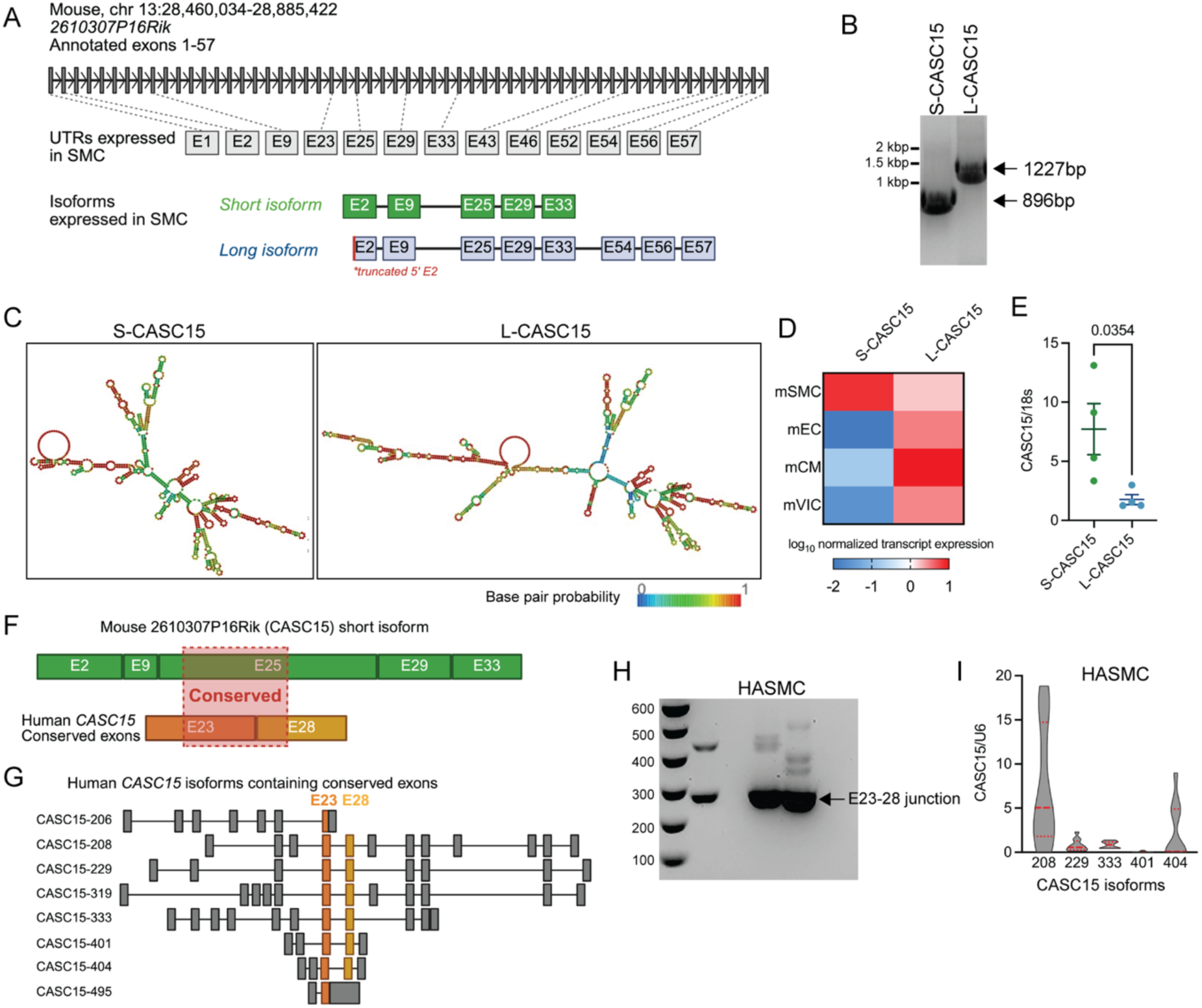
Vascular SMCs express a conserved lineage-selective CASC15 isoform. **A**. Schematic presentation of *Mus musculus 2610307P16Rik* and transcripts sequenced in mSMCs. **B**. PCR amplification of full-length S-CASC15 and L-CASC15 transcripts in mSMCs. **C**. Modeling of S-CASC15 and L-CASC15 conformations using Vienna Suite. **D**. S-CASC15 and L-CASC15 expression in mouse SMC, endothelial cells (mEC), cardiomyocytes (mCM), and valve interstitial cells (mVIC). CASC15 expression was normalized to 18s. **E**. Comparative expression of S-CASC15 and L-CASC15 in mSMC. Student t-test. **F**. Schematic representation of sequence homology and conservation between mouse S-CASC15 and human CASC15. The junction between human CASC15 E23 and E28 untranslated exons exhibits high conservation relative to S-CASC15. **G.** Representation of annotated CASC15 transcript isoforms containing the untranslated exons E23 and E28. **H.** PCR amplification of the E23-28 junction in HASMC. **I.** Quantification of human CASC15 transcripts containing the E23-28 junction. Expression normalized to U6.

### CASC15 deficiency promotes SMC hypertrophy and polyploidization

We evaluated the function of CASC15 in mouse aortic SMCs by designing isoform-specific and non-selective LNA antisense oligonucleotide GapmeRs (**Figure 3A**). CASC15 GapmeRs induced a marked reduction in CASC15 levels compared to control GapmeR, with no significant difference between CASC15 GapmeR #1 (S-CASC15) or #2 (non-selective), supporting the dominant role of S-CASC15 in SMCs (**Figure 3B**). CASC15 knockdown (KD) resulted in notable cell hypertrophy (Figure 3C). Further assessment of morphological features revealed increases in cell and nuclear size, as well as in the percentage of binucleated cells (**Figure 3D**). Importantly, cellular hypertrophy was selectively induced by S-CASC15, whereas no effects were observed when silencing L-CASC15 (**Supplementary Figure 7A**). Hypertrophy and ploidy were confirmed with electron microscopy, with the presence of enlarged and binucleated SMCs following CASC15 KD (**Figure 3E**, **Supplementary Figure 7B**). Beyond the occurrence of binucleation, flow cytometry revealed that CASC15 KD led to a marked increase in the percentage of polyploid (>2N) cells, reaching 38.6% in CASC15 KD SMCs (**Supplementary Figure 7C**). These observations were recapitulated in HASMCs, in which CASC15 KD also triggered cell hypertrophy (**Figure 3 F-G**). Aortic SMC hypertrophy and polyploidy are key features reported in hypertension and vascular aging^13,14,63,64^. Mechanistically, angiotensin II (ANG-II) has been identified as a key driver of SMC hypertrophy and polyploidization^14,65^. Treatment with ANGII for 24 h led to a robust reduction in CASC15 expression in SMC across species and vascular territories (**Figure 3H, Supplementary Figure 8A**). These observations were recapitulated in vivo, following 28 days of ANGII infusion in mice, resulting in decreased CASC15 levels in the aorta and renal artery (**Figure 3I**). Treatment with Olmesartan, a selective Angiotensin II Receptor type 1 inhibitor, prevented CASC15 level reduction (**Supplementary Figure 8B**). CASC15 KD had a comparable effect on SMC cell and nucleus size to ANGII in serum-starved SMCs (Figure 3J-K). The combination of CASC15 KD and ANGII treatment did not exhibit a cumulative or synergistic effect, supporting the notion that ANGII effects are CASC15-dependent. Furthermore, CASC15 overexpression significantly rescued ANGII-mediated SMC hypertrophy (**Figure 3L-M**). Of note, CASC15 overexpression did not significantly change cell size in SMCs treated with vehicle. These studies uncover the critical role of CASC15 in regulating SMC cell size. Downregulation of CASC15 by ANGII is a determinant step driving SMC hypertrophy, binucleation, and polyploidization.

**Figure 3:**
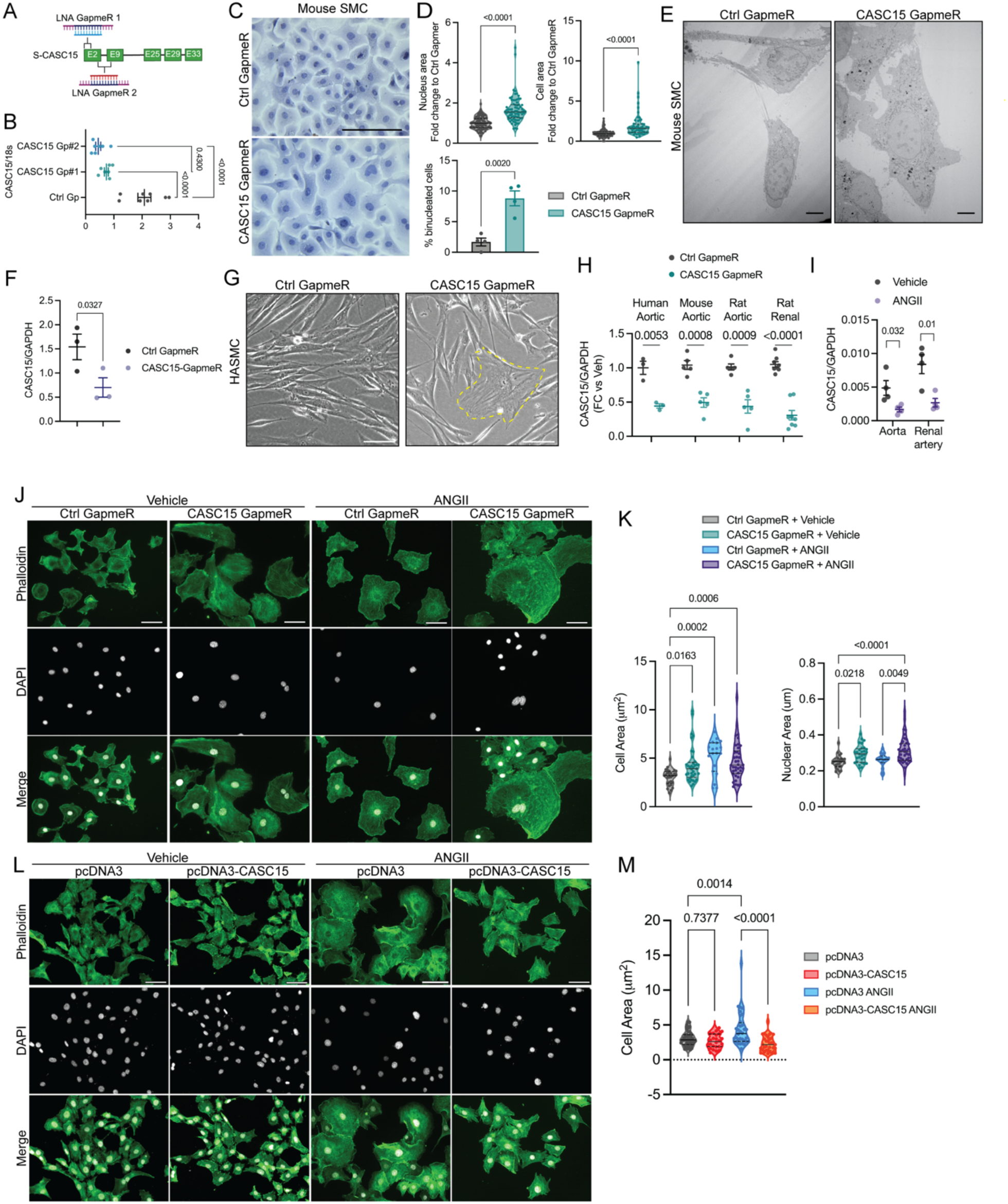
CASC15 deficiency drives SMC hypertrophy and polyploidization. **A**. LNA GapmeRs were designed for selective (S-CASC15, CASC15 GapmeR #1) or non-selective (CASC15 GapmeR #2) targeting of CASC15. **B.** CASC15 expression in mSMC treated with control and CASC15 Gapmers. Data expressed as mean ± SEM. One-way ANOVA. **C.** Representative micrographs of control and CASC15 GapmeR-treated mSMC. Scale bar = 50 µm. **D.** Cell size, nucleus size, and percentage polynucleation rate in control vs CASC15 GapmeR. Data expressed as mean ± SEM. Student’s t-test. **E**. Electron microscopy representative images of SMC after transfection with control vs CASC15 GapmeR. Scale bar = 1 µm. **F.** Gapmer-mediated CASC15 knockdown in HASMC. CASC15 expression normalized to GAPDH. Data expressed as mean ± SEM. Student t-test. **G.** Representative micrographs of control and CASC15 KD HASMC. Scale bar = 50 µm. **H**. CASC15 expression in ANGII-treated (100nM, 24h) SMCs. CASC15 expression was assessed in aortic human, mouse, and rat SMCs, as well as rat renal SMCs. Expression normalized to GAPDH. Data expressed as mean ± SEM. Multiple Student’s t-test. **I.** CASC15 expression in the aorta and renal artery of ANGII and vehicle-infused mice (1000 ng/kg/min). **J**. Representative images of control and CASC15 GapmeR SMC treated with vehicle or ANGII for 24 hours. Staining: Phalloidin (green) and DAPI (gray). Scale bar = 50 µm. **K**. Quantification of cell and nucleus size. Data expressed as mean ± SEM. One-way ANOVA. **L**. Representative images of control and CASC15-overexpressing SMC treated with vehicle or ANGII for 24 hours. Staining: Phalloidin (green) and DAPI (gray). Scale bar = 50 µm. **M**. Quantification of cell size. Data expressed as mean ± SEM. One-way ANOVA.

### CASC15 drives SMC proliferation and ensures cell cycle fidelity

Mouse aortic SMCs with reduced CASC15 expression experienced a decrease in proliferating cells compared to their control counterpart as measured by BrdU incorporation and cell count (**Figure 4A and 4C**, **Supplementary Figure 9**). CASC15 knockdown also abolished the mitogenic potential of Platelet-Derived Growth Factor-BB (PDGF-BB), known to potently drive SMC proliferation^66^. Interestingly, CASC15 overexpression had no effect on SMC viability or proliferation in the absence of growth factors (**Figure 4C-D, Supplementary Figure 9**). However, proliferation of CASC15-overexpressing SMCs was enhanced in the presence of PDGF-BB. A similar pro-proliferation effect of CASC15 was observed in HASMCs (**Figure 4D**). Collectively, these data support a role for CASC15 in enhancing SMC proliferative capacity, particularly under conditions that promote cell-cycle entry. Consistently, we found an increase in the proportion of CASC15-deficient SMCs in the G1 phase and a concomitant decrease in the proportion of cells in the G2/M phase (**Figure 4E**). Interestingly, in proliferating SMCs, CASC15 exhibited cyclic expression, peaking at 6 hours after PDGF-BB treatment, then returning to baseline at 12 hours (**Figure 4F**). To further evaluate the role of CASC15 in regulating cell cycle, we transduced SMC with the Fluorescent Ubiquitination-based Cell Cycle Indicator (FUCCI) reporter, which differentially labels cells in G1 (red), S (yellow), and G2/M (green) (**Figure 4G**)^67^. FUCCI-SMCs cultured with serum-containing media were flow-sorted to isolate cells in G1, S, and G2/M phases. Consistent with the PDGF-BB time course, CASC15 expression fluctuated during the cell cycle, with the highest levels observed in G2/M-phase SMCs (**Figure 4H**). Live imaging using FUCCI-SMCs transfected with CASC15- or control GapmeR revealed that CASC15 depletion was associated with an accumulation of polyploid, binucleated SMC in G1 (**Figure 4I, Supplementary Figure 10**). These results suggest that CASC15 expression during the G2/M phase is essential for cell cycle completion and fidelity, and that its absence leads to a G2/M phase defect, incomplete mitosis, and subsequent G1 phase arrest. To further evaluate how CASC15 regulates SMC functions and phenotypes, we performed bulk RNA-seq on CASC15-deficient SMCs treated with vehicle or PDGF-BB, CASC15-overexpressing SMCs, and their respective controls (**Supplementary Figure 11A, Supplementary Tables 5-8**). All treatments were associated with marked changes in gene expression as compared to control SMCs (**Supplementary Figure 11B**). The most differentially enriched pathways were associated with immune responses (**Supplementary Figure 11C-D**). These results align with recent findings suggesting that CASC15 promotes endothelial cell inflammation in pro-atherogenic in vitro models^58^. Beyond inflammatory response, pathways associated with the cell cycle were differentially enriched in CASC15-deficient cells (**Figure 4J, Supplementary Figure 12**). Specifically, G1/S and G2/M phase transition regulation, as well as cytokinesis pathways, were downregulated upon CASC15 KD, whereas regulation of cell growth and mitotic cell cycle arrest pathways was upregulated. Gene Set Enrichment Analysis (GSEA) showed a significant reduction in the expression of genes associated with metaphase-anaphase transition (**Figure 4K-L**). We found a transcriptional signature consistent with the FUCCI imaging, comprising the upregulation of genes involved in G1 arrest (Cdkn1a, Pidd1, and Cdc73), downregulation of genes that prevent G1 arrest (Gli1), and downregulation of G2/M transition checkpoint genes (Chek2, Cenpt, Eme1, Cit) (**Figure 4M**). Conversely, CASC15 overexpression was associated with upregulation of cell cycle pathways (**Figure 4N**). Together, these studies demonstrate that CASC15 plays a central role in ensuring cell cycle fidelity. Deficiency in CASC15 expression in SMCs that reenter the cell cycle leads to incomplete mitosis and subsequent G1 cell cycle arrest. Meanwhile, increased CASC15 expression exacerbates SMC proliferation. These observations interrogate the consequences of variation in CASC15 expression during vascular remodeling in vivo. In addition to its role in regulating the cell cycle, we found that CASC15 alterations profoundly alter the expression of multiple extracellular matrix components, potentially impacting vascular remodeling (**Supplementary Figure 13**).

**Figure 4:**
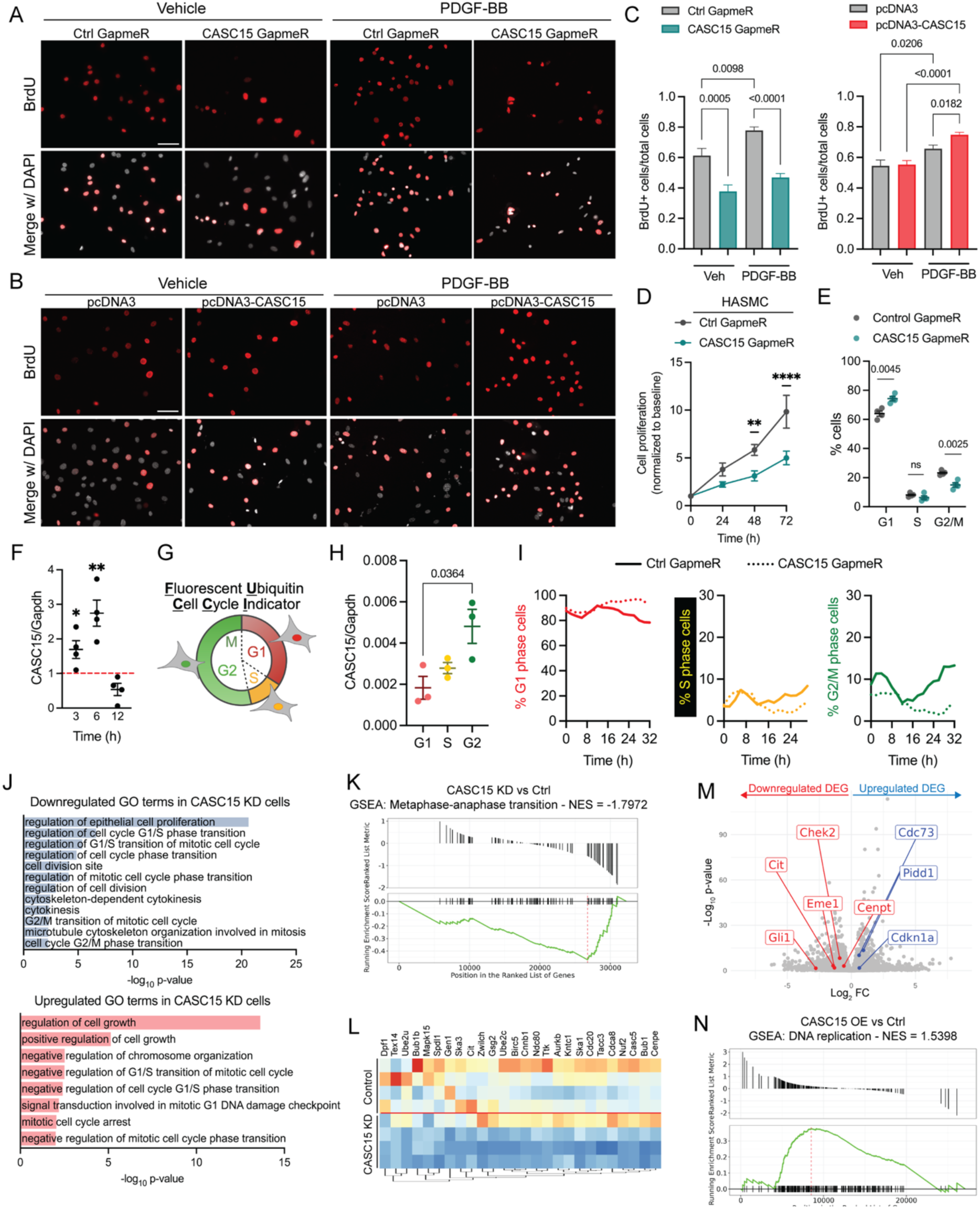
CASC15 warrants G2/M transition, mitosis transition, and cell cycle fidelity. **A**. Representative micrographs of BrdU incorporation in control and CASC15-deficient mSMC treated with PDGF-BB (10 ng/mL, 24 hours) or vehicle. Nuclei counterstained with DAPI. Scale bar = 50 µm. **B**. Representative micrographs of BrdU incorporation in control and CASC15-overexpressing mSMC treated with PDGF-BB (10 ng/mL, 24 hours) or vehicle. Nuclei counterstained with DAPI. Scale bar = 50 µm. **C.** BrdU+ cell quantification. Data expressed as mean ± SEM. One-way ANOVA. **D.** Cell growth over time in HASMC transfected with Control or CASC15 GapmeR. Data expressed as mean ± SEM. Two-way ANOVA. ** p < 0.001, **** p < 0.0001. **E.** Propidium iodide flow cytometry analysis of cell cycle phases in Control- and CASC15-GapmeR mSMC. Data expressed as mean ± SEM. Multiple t-test. **F.** CASC15 expression in mSMC during PDGF-BB (10 ng/mL) time course. Results are expressed as fold change compared to baseline (0h). Data expressed as mean ± SEM. Multiple t-test. **G.** Schematics of the FUCCI reporter system. **H.** CASC15 expression in FUCCI-sorted cells. Data expressed as mean ± SEM. One-way ANOVA. **I.** Quantification of the percentage of SMC in G1 (red), S (yellow), and G2 (green) phases over time after transfection with Control- or CASC15-GapmeR. **J.** Gene Ontology analysis of significantly downregulated or upregulated cell cycle-related pathways in CASC15-deficient SMC. **K.** Metaphase-anaphase transition Gene Set Enrichment Analysis in CASC15-deficient SMC vs control. **L.** Heat map of the metaphase-anaphase transition gene set. **M.** Volcano plot showing significant differential expression of genes involved in G1 arrest (Cdc73, Pidd1, Cdkn1a, Gli1) and G2/M checkpoint (Chek2, Cit, Eme1, Cenpt) in CASC15-deficient SMC vs control. **N.** DNA replication Gene Set Enrichment Analysis in CASC15-overexpressing SMC vs control.

### CASC15 drives hyperplastic vascular remodeling by promoting cell proliferation and migration

The modulation of SMC behavior, whether by hyperplastic or hypertrophic growth, has been shown to be a major contributor to vascular remodeling in many diseases^68^. We tested the role of CASC15 in vascular remodeling through a series of loss- and gain-of-function studies in a ligation-induced carotid artery model (**Figure 5A**). Although this model does not recapitulate the complex pathogenesis of vascular diseases such as atherosclerosis, its hyperplastic nature makes it highly relevant for studying the impact of CASC15 modulation on SMC proliferation in vivo. We performed unilateral right carotid artery ligation and, concomitantly, delivered Control- or CASC15-shRNA lentivirus into SMC fate-mapping mice. CASC15 KD induced an almost complete absence of neointima formation (**Figure 5B-C**). However, CASC15 KD was associated with an increase in the medial-to-vessel area ratio, suggestive of hypertrophic remodeling. This was confirmed by quantification of cell density, which was lower in CASC15 KD arteries, indicating that the increase in medial area is not due to an increase in SMC number, but to an increase in their mass (**Figure 5C**). These observations aligned with the in vitro studies and suggested that the lack of CASC15 markedly impairs cell proliferation and promotes cell hypertrophy. In addition to its role in controlling cell proliferation, we found that loss of CASC15 impeded SMC migration (**Figure 5D**, **Supplementary Figure 14**). Consistent with these observations, RNA sequencing revealed profound alterations in the expression of genes associated with taxis and migration (**Figure 5E**). CASC15 overexpression increased hyperplasia and neointima lesion size (**Figure 5F-G**). The increase in neointima formation was also associated with a reduction in the media-to-vessel ratio as compared to control animals. Importantly, the increase in neointima area was driven by an increase in the number of SMCs (fate-mapping reporter positive). To further evaluate the role of CASC15 in pathological vascular remodeling and based on scRNAseq and scATACseq data showing increased expression in fibromyocytes (**Supplementary Figure 3**), we generated a novel CASC15 knockout (KO) mouse model and induced atherosclerosis through a combination of PCSK9 AAV delivery and high-fat diet. CASC15 KO mice were generated by CRISPR-Cas9-mediated excision of the *CASC15* proximal promoter, untranslated first exon, and first intron (**Supplementary Figure 15A-B**). This manipulation resulted in a complete loss of CASC15 in all organs tested, including the aorta (**Supplementary Figure 15 C-D**). CASC15 KO mice were viable and maintained a similar body weight to wild-type littermates (**Supplementary Figure 15E**). CASC15 deletion was not associated with gross organ morphological abnormalities (**Supplementary Figure 15F**). CASC15 KO mice were subjected to tail vein injection with PCSK9-AAV, followed by 16 weeks of high-fat diet feeding. CASC15 KO mice exhibited smaller atherosclerotic lesions in the brachiocephalic artery than their wild-type littermates (**Figure 5H-I**). Atherosclerotic lesions in CASC15 KO mice were rich in macrophages relative to ACTA2+ cells, which suggests a possible impairment of medial SMC proliferation and investment of the lesion (**Figure 5J**). Together, our studies provide evidence that CASC15 plays an essential role in promoting lesion expansion after vascular injury and during atherosclerosis.

**Figure 5:**
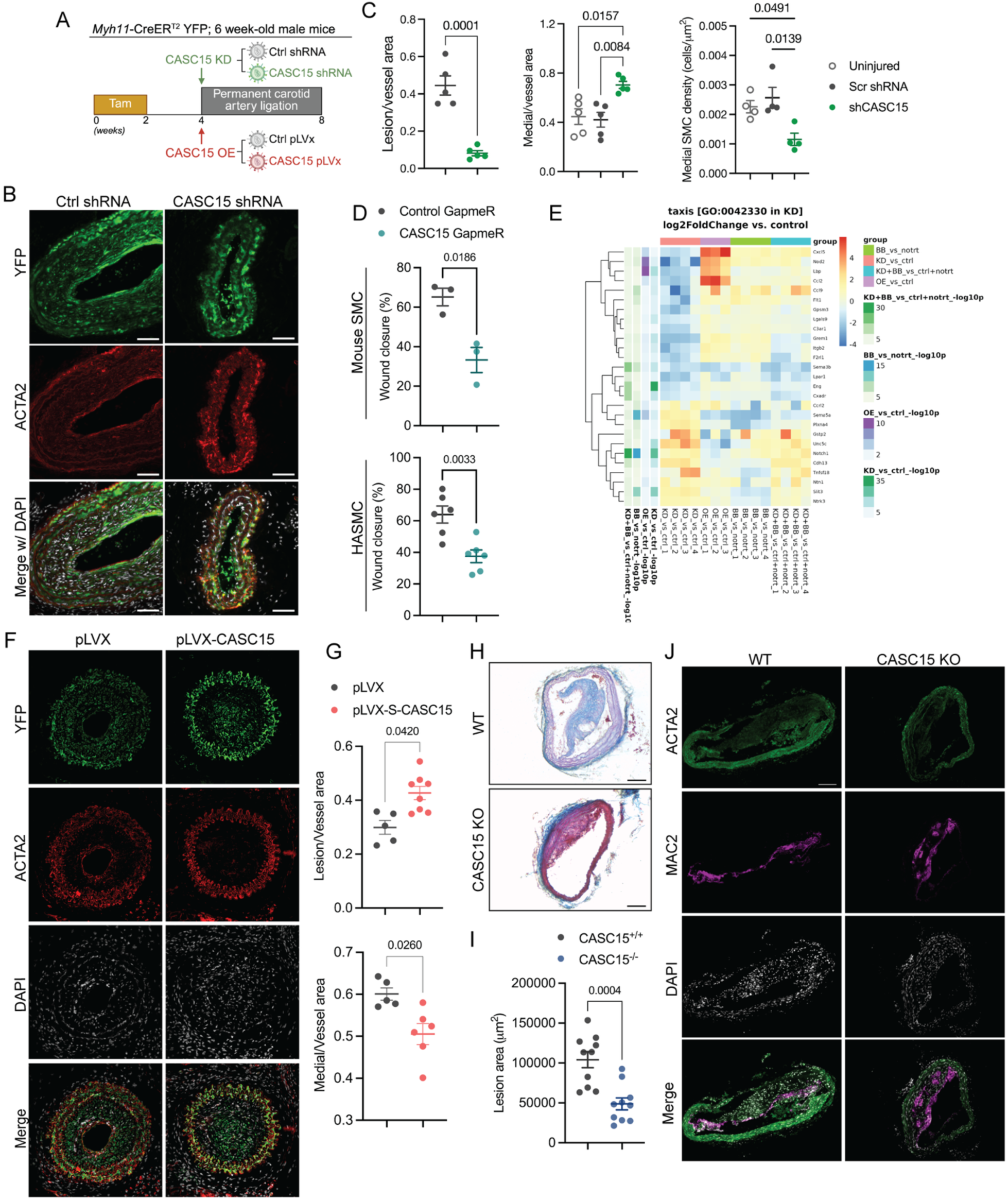
CASC15 promotes hyperplastic vascular remodeling. **A.** Schematic representation of permanent ligation and simultaneous delivery of Pluronic gel containing CASC15/Control shRNA lentivirus into the right carotid artery of *Myh11*-CreER^T2^ YFP mice. Mice were injected with tamoxifen two weeks before procedures for definitive SMC labeling**. B.** Representative micrographs of ligated carotid arteries stained with YFP (SMC reporter), ACTA2, and DAPI. Scale bar = 100 μm. **C.** Morphometric analysis of lesion/vessel area, medial/vessel area, and medial SMC density (number of cells/μm^2^). Data expressed as mean ± SEM. Student t-test and One-way ANOVA. **D.** Quantification of mouse (top) and human (bottom) SMC migration in scratch assay. Data expressed as mean ± SEM. Student t-test. **E.** Heat map of genes related to taxis and migration in experimental groups vs controls. FC: Fold change. **F.** Representative micrographs of ligated carotid arteries receiving pLVX-CASC15 or control virus and stained with YFP (VSMC reporter), ACTA2 antibodies, and DAPI. Scale bar = 100 μm. **G.** Morphometric analysis of lesion/vessel area and medial/vessel area. Data expressed as mean ± SEM. Student t-test. **H.** Representative micrographs of brachiocephalic arteries harvested from PCSK9-AVV-injected and 16-week high-fat diet-fed CASC15^-/-^ mice and WT littermates. Scale bar = 100 μm. **I.** Morphometric analysis of brachiocephalic artery atherosclerotic lesion area in CASC15^-/-^ mice and WT littermates. Data expressed as mean ± SEM. Student t-test. **J.** Immunofluorescent staining with ACTA2, MAC2 antibodies, and DAPI. Scale bar = 100 μm.

### CASC15 deficiency induces spontaneous medial hypertrophy and vessel hypercontractility

Our in vitro data demonstrate that reduced CASC15 expression drives SMC hypertrophy (**Figure 3**). Similarly, in vivo delivery of CASC15-shRNA to injured carotid arteries resulted in medial hypertrophy and a lack of neointima formation (**Figure 5**). We further investigated the morphological and functional changes in the vasculature associated with constitutive global CASC15 KO. CASC15 KO mice exhibited spontaneous aortic hypertrophy, characterized by increased medial area and thickness (**Figure 6A-B**). Wheat Germ Agglutinin (WGA) staining was performed to precisely assess medial SMC morphology. Deletion of CASC15 resulted in a marked increase in cell size but not cell number, demonstrating that the medial hypertrophy is due to increased SMC mass rather than hyperplasia (**Figure 6C-D**). Consistent with our in vitro observations, we also found an increase in SMC nucleus size and a higher frequency of binucleation in the aorta of CASC15 KO mice compared to their control littermates (**Figure 6D**, **Supplementary Figure 16A**). Interrogation of the RNA-seq data and GO pathway analysis performed on CACS15 KD SMC revealed upregulation of pathways related to SMC contractility and dysregulation of pathways associated with vasoreactivity and blood pressure regulation (**Figure 6E**). Increased ACTA2 expression was found in the aorta of CASC15 KO mice as well as in CASC15-deficient SMC (**Supplementary 16B-C**). Importantly, medial hypertrophy was also detected in mesenteric resistance arteries (**Figure 6F**). Vessel thickening (i.e., increase in the wall-to-lumen ratio) has been shown to elevate vascular resistance and reduce arterial compliance to blood vessel tone^63,69^. We performed *ex vivo* vessel wire myography experiments using CASC15 KO and WT mesenteric arteries and aortas to assess their constriction capacities in response to phenylephrine and U46619 (a stable thromboxane A_2_ mimetic) (**Figure 6G**). CASC15 KO mesenteric arteries exhibited higher contractility in response to both vasoconstrictors (**Figure 6H**). Finally, CASC15 KO mesenteric arteries and aortas display enhanced KCl-induced maximal contraction compared with WT mice (**Figure 6I**). In summary, constitutive loss of CASC15 induces major spontaneous morphological and functional alterations of the vasculature, including medial and SMC hypertrophy and SMC hypercontractility, as well as resistance and conductance artery hyperconstriction.

**Figure 6:**
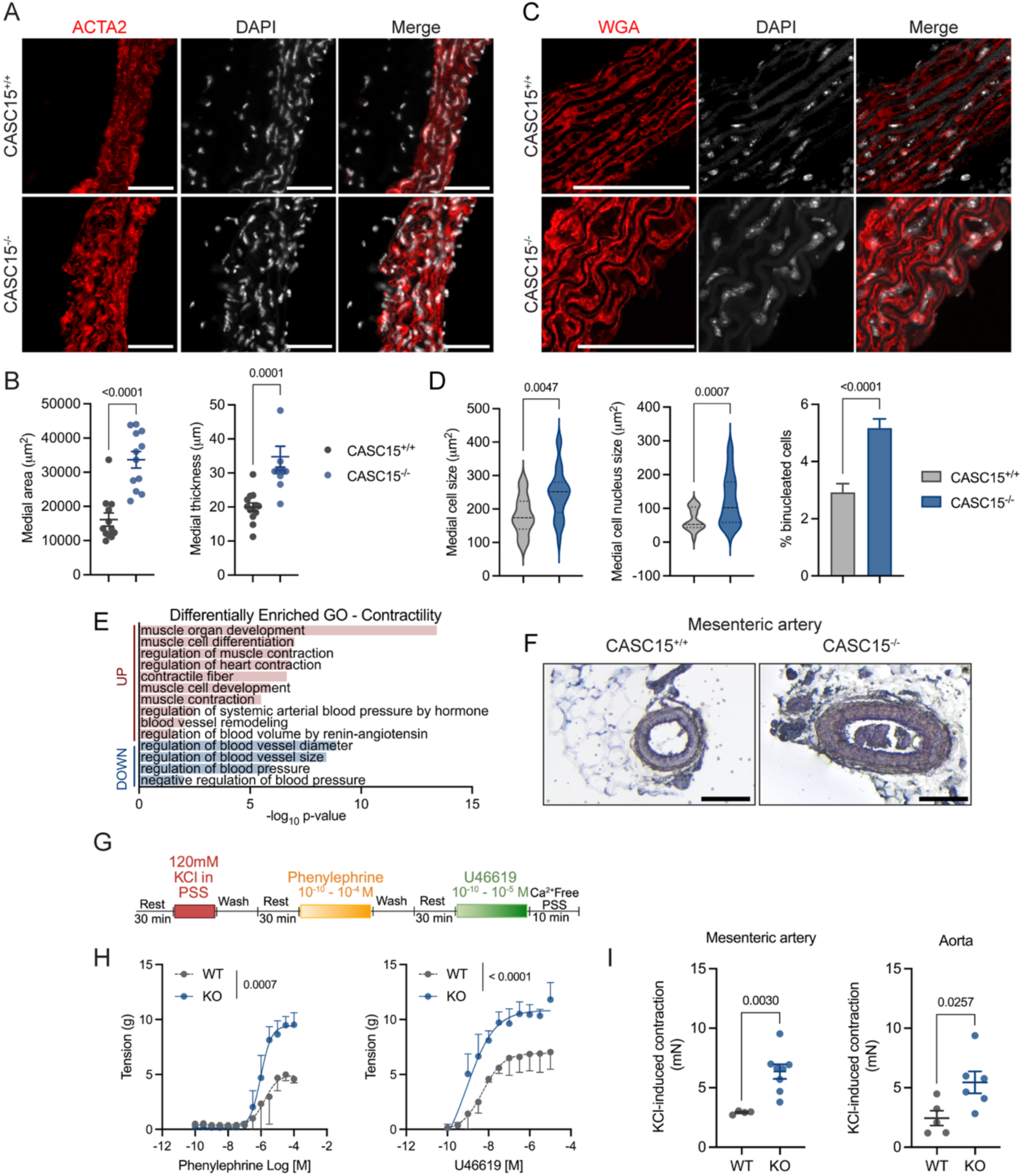
CASC15 deficiency induces vascular hypertrophy and hypercontractility. **A.** Representative micrographs of CASC15^+/+^ and CASC15^-/-^ aortas stained for ACTA2 and DAPI. Scale bar = 25μm. **B.** Morphometric analysis: CASC15^-/-^ and CASC15^+/+^ aorta medial area and thickness. Data expressed as mean ± SEM. Student t-test. **C.** Representative micrographs of WGA and DAPI staining in CASC15^+/+^ and CASC15^-/-^ aortas. Scale bar = 25μm. **D.** Quantification of cell size, nucleus size, and binucleation frequency. Data expressed as mean ± SEM. Student t-test. **E.** Gene ontology analysis uncovers significantly differentially regulated pathways associated with contractility, blood vessel remodeling, vasoreactivity, and blood pressure. Upregulated pathways are highlighted in red, whereas downregulated pathways are highlighted in blue. **F.** Representative micrographs of mesenteric arteries stained with Masson. Scale bar = 100μm. **G.** Schematics of vascular reactivity assessment by ex vivo wire myography. **H.** Mesenteric artery contractility in response to phenylephrine and U46619 dose response. Data expressed as mean ± SEM. Two-way ANOVA. **I.** KCl-induced maximal constriction of CASC15^+/+^ and CASC15^-/-^ mesenteric arteries and aortas. Data expressed as mean ± SEM. Student t-test.

### CASC15 regulates cell cycle fidelity and cell morphology by controlling RNA stability

We performed RNA pull-down followed by mass spectrometry to identify putative CASC15-interacting partners and mechanisms of action (**Figure 7A**). A clear separation was observed between protein extracts incubated with biotinylated-S-CASC15 probe and those incubated with the biotinylated-CASC15 AntiSense control probe (**Supplementary Figure 17A**). Raw Data-Independent Acquisition (DIA) proteomics data were processed with FragPipe and Spectronaut, which identified 73 and 47 significantly differentially enriched interacting candidates in biotinylated-CASC15 samples based on the following cutoffs: q-value = 0.05, Log_2_ fold change ≥ 0.58 (**Supplementary Tables 9-12**, **Supplementary Figure 17B**). Thirty-nine (39) overlapping targets were commonly reported as significantly enriched in biotinylated-S-CASC15 samples (**Figure 7B**). String and gene ontology analysis revealed that most of the 39 candidates were nuclear proteins involved in RNA processing (i.e. RNA splicing, stability) and translation initiation (**Figure 7C-D**). Consistent with these findings, we observed that CASC15 exhibits nuclear and perinuclear localization and thus resides in the same subcellular compartments as the pulldown candidates (**Supplementary Figure 18**). Among the identified candidates were multiple small nuclear ribonucleoproteins (SNRNPs), ribosomal proteins, and the multifunctional RNA-binding protein Nucleolin (Ncl), suggesting that CASC15 participates in the assembly or function of ribonucleoprotein complexes. RNA-protein interaction simulation using AlphaFold3 predicted that CASC15 forms a complex with SNRNP70, as well as Nucleolin in its unphosphorylated and phosphorylated states (**Figure 7E, Supplementary Figure 19A, Supplementary Video 1-3**). Furthermore, Ncl and SNRNP70 can bind CASC15 at the same time to form a ternary complex (**Supplementary Figure 19B**, **Supplementary Video 4**). We then postulated that CASC15-dependent regulation of transcript stability may contribute to the shift from proliferative toward hypertrophic growth and cell-cycle arrest. RNA stability assays revealed that CASC15 deficiency was associated with shifts in transcript stability of cell cycle-associated proteins. Specifically, CASC15 KD decreased the stability of Chek2, a G2/M checkpoint protein (**Figure 7F**). In contrast, the stability of Cdkn1a (associated with G1 arrest) and Cks1 (associated with SMC binucleation^70^) was enhanced upon CASC15 KD. These results suggest that dysregulation of cell cycle proteins, characterized by downregulation of G2/M checkpoint proteins and upregulation of G1 arrest-associated proteins, could be associated with impairment of CASC15-dependent selective transcript stability (**Figure 4**). Ncl regulates not only central RNA metabolism processes (RNA splicing and stability^71^) but has also been implicated in cell cycle control. Indeed, studies have shown that Ncl depletion induces cell binucleation^72^. Performing siRNA-mediated Ncl knockdown, we observed that the reduction in Ncl phenocopied CASC15 KD, with evident SMC hypertrophy, binucleation, and impaired cell proliferation (**Figure 7G-J**). Of major importance, the overexpression of S-CASC15 in Ncl-deficient SMC did not rescue these cellular features, indicating that the effects of CASC15 are Ncl-dependent. These mechanistic studies provide compelling evidence that CASC15 regulates cell cycle and cell morphology by controlling RNA processing and stability in an Ncl-dependent fashion. Our data also suggest that CASC15 serves as a scaffold molecule for the formation of specific ribonucleoprotein complexes (e.g. Snrnp70 and Ncl) that selectively regulate the stability of a set of transcripts involved in cell cycle completion. Together, CASC15 is a central mechanism regulating vascular SMC growth fate, and whose expression level biases vascular remodeling toward SMC hyperplasia or hypertrophy.

**Figure 7:**
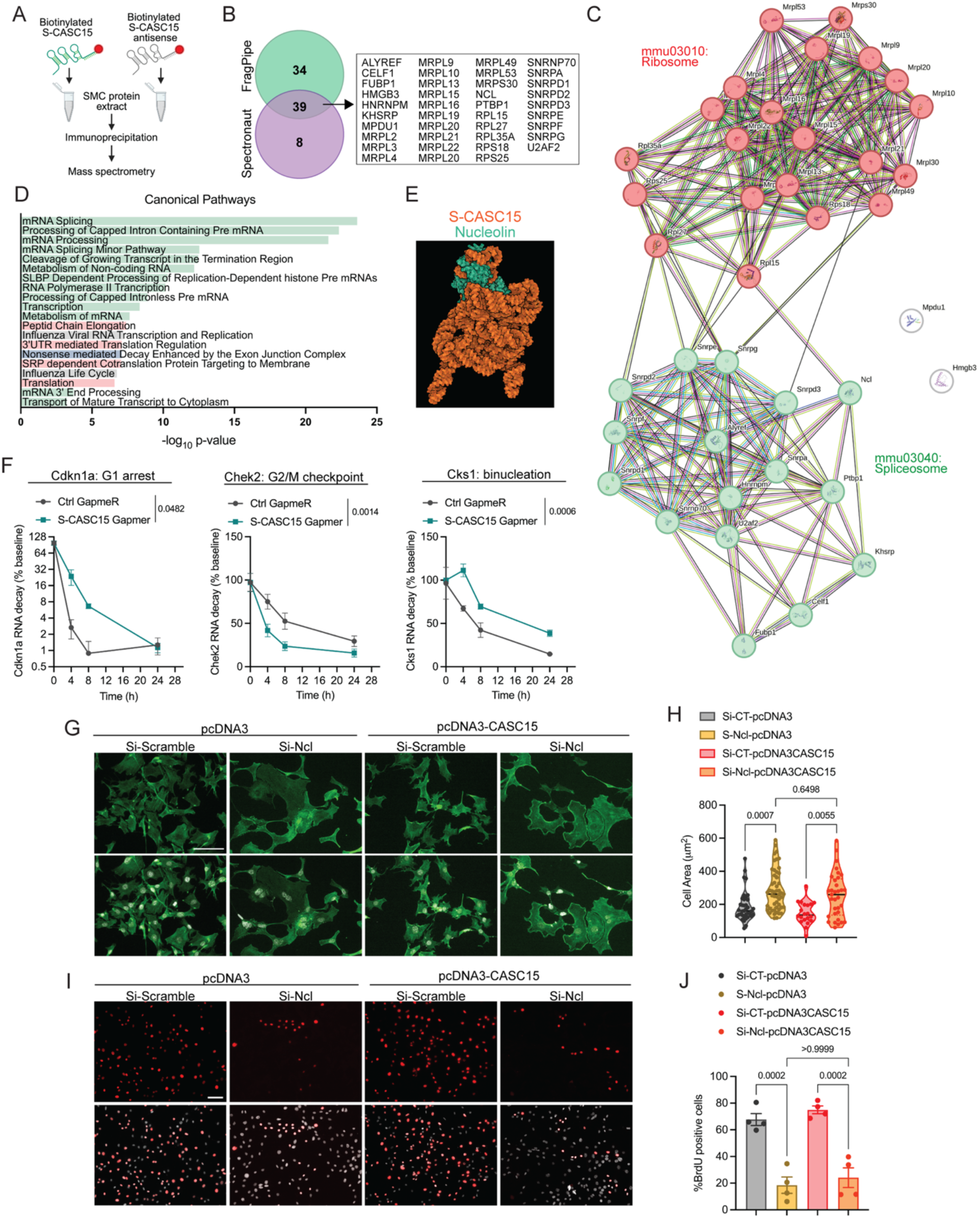
CASC15 regulates cell division and morphology through control of RNA stability. **A**. Schematics of the RNA/Protein pulldown using biotinylated probes and followed by mass spectrometry. **B.** CASC15 interacting partners candidates identified by Fragpipe and Spectronaut. **C.** String analysis of the 39 common candidates revealed enrichment in proteins involved in RNA splicing and translation. **D.** Gene Ontology analysis based on the list of 39 common candidates. **E.** AlphaFold3 simulation predicting S-CASC15 and Nucleolin interaction. **F.** RNA decay assay in CASC15- and Control-GapmeR SMCs treated with Actinomycin D. Cdkn1a, Check2, and Cks1 transcript decay was quantified over time. Data expressed as mean ± SEM. Two-way ANOVA. **G.** Representative micrographs of SMCs transfected with siNcl or control and pcDNA3-CASC15 or control. Staining: Phalloidin (green), DAPI (gray). Scale bar = 50 μm. **H.** Quantification of cell area in experimental groups outlined in G. Data expressed as mean ± SEM. One-way ANOVA. **I.** Representative images of SMC proliferation by BrdU incorporation. Scale bar = 50 µm. **J.** Quantification of SMC proliferation based on BrdU positivity. Data expressed as mean ± SEM. One-way ANOVA.

## DISCUSSION

Our study identifies CASC15 as a key lncRNA that regulates vascular remodeling by modulating SMC morphology and cell cycle completion. We found a novel SMC-selective CASC15 isoform that is tightly regulated by growth factors and Angiotensin-II. Expression of CASC15 is essential for mitosis completion and cell cycle fidelity. Variations in CASC15 expression play a causal role in regulating SMC growth fate during in vivo vascular remodeling: while an increase in CASC15 expression in G2/M phase cells promotes SMC hyperplasia, CASC15 deficiency drives SMC and vascular hypertrophy and hypercontractility. Collectively, these findings position CASC15 as a molecular gateway governing the nature of SMC participation in vascular function, remodeling, and disease progression.

While the role of CASC15 in regulating cell proliferation and growth is well documented across various cancers, its precise expression profile and functional impact remain controversial ^49,52,54,57,73,74^. This ambiguity likely stems from its extensive isoform diversity, which complicates the investigation of CASC15 function but also helps explain the heterogeneity and possible tissue-specificity of the effects and mechanisms reported to date. The Ensembl Genome Browser (v115) identifies approximately 300 human and 242 mouse transcripts. For instance, while one CASC15 isoform promotes neuroblastoma, another transcript was observed to suppress tumor growth, with lower levels associated with worse patient prognosis^75^ ^73^. In vascular SMC, we detected two major CASC15 transcripts: a short and a long isoform (S-CASC15 and L-CASC15, respectively). Importantly, these isoforms differ from previously annotated transcripts. Whereas L-CASC15 is detected in many cardiovascular cell types and highly expressed in cardiomyocytes, S-CASC15 is remarkably selectively expressed in vascular SMC. While our initial focus on S-CASC15 was motivated by its selective expression in SMC, in vitro and in vivo loss-of-function studies demonstrated that S-CASC15 is the functionally dominant isoform in SMC. In apparent contradiction to its pro-proliferation effects in SMC, previous studies reported that CASC15 overexpression induces cardiomyocyte hypertrophy^57^. This discrepancy may be explained by the expression of different CASC15 isoforms in cardiomyocytes and SMCs, and by their possible divergent effects on cell cycle regulation and cell morphology. Future studies should further delineate the context-and cell-type-dependent expression profile of CASC15 transcripts and evaluate the function of the identified isoforms.

CASC15 is expressed in quiescent SMCs and is selectively enriched in human vascular and visceral tissues. Our study revealed that variation in CASC15 expression levels dictates the mode of SMC growth during vascular remodeling: high CASC15 expression promotes hyperplasia, while SMCs undergo hypertrophy when CASC15 is downregulated. Interestingly, CASC15 expression varies during the cell cycle, with significant enrichment during the G2/M phase. These findings make a compelling case for considering CASC15 a “cycling lncRNA”. Other lncRNAs exhibit similar features. Hao et al. unbiasedly characterized lncRNA expression in cell-cycle-synchronized osteosarcoma cells and found that many lncRNAs display differential expression across distinct phases of the cell cycle^76^. In the study’s dataset, CASC15 expression is higher in the S and G2 phases than in the G1 and M phases. MALAT1 and GAS5, two lncRNAs implicated in regulating SMC phenotypic states, also exhibit cell-cycle phase-dependent mechanisms^36,77,78^. In vitro knockdown and in vivo knockout of CASC15 resulted in a high rate of SMC polyploidy and binucleation. Transcriptomic analysis revealed that loss of CASC15 expression led to downregulation of G2/M checkpoint proteins and increased expression of G1 arrest-associated genes. Together, our data support the hypothesis that the absence of CASC15 impairs proper mitotic completion, leading to subsequent G1 cell arrest. Our observations align with a body of research linking mitosis failure and tetraploidy to p53-mediated G1 arrest^79,80^. We also found that CASC15 KD led to the overexpression of Cks1. Importantly, Cks1 expression is increased in the media of hypertensive rat aortas, as well as in SMC treated with ANGII^70^. Hixon et al. also showed that overexpression of Cks1 in aortic SMC from normotensive rats was sufficient to cause polyploidization. Binucleation, polyploidy, and mitosis failure can arise through several distinct mechanisms. Cells can exit mitosis through mitotic slippage, cytokinesis failure (also referred to as acytokinetic mitosis), or endoreplication. While mitotic slippage and endoreplication generally generate polyploid cells with a single nucleus, cytokinesis failure triggers the formation of binucleated cells^81–83^. Whether CASC15 deficiency induces one or several of these mitosis defect mechanisms remains to be investigated.

Modulation of CASC15 expression resulted in striking CVD-relevant phenotypes. Firstly, CASC15 overexpression led to neointimal expansion driven by an increase in the number of SMCs. Both proliferation and migration were positively regulated by CASC15. Consistent with these observations, CASC15 KD and KO caused defects in vascular injury-induced neointimal formation and reduced atherosclerotic lesion size. Importantly, the decrease in lesion area appeared to be driven by reduced SMC investment, a lack of ACTA2^+^ cell-containing fibrous cap formation, and macrophage accumulation. These results interrogate the overall effect of CASC15 deficiency on atherosclerosis formation and progression, as plaque size itself is considered a poorer predictor of plaque rupture risk than indices of plaque stability, particularly fibrous cap thickness^84–86^. Given the potential significance of CAS15 in regulating atherosclerosis pathogenesis, two questions remain to be investigated. Firstly, the marked effect of CASC15 on proliferation interrogates its involvement in SMC clonality. Murine preclinical and human studies have revealed that SMCs propagate and populate atherosclerotic lesions through extensive proliferation of a limited number of medial SMCs^11,12,87^. While the underlying mechanisms remain to be further investigated, SMC clonality can arise from the proliferative propensity of a primed SMC subpopulation and/or environmental clonal selection^88^. It has been postulated that a small population of medial Sca1+ SMC clonally expands and populates atherosclerotic lesions^11,89^. Similarly, restricting CASC15 cyclic upregulation to a small number of SMCs may mediate oligoclonal expansion, whereas indiscriminate overexpression would favor polyclonal proliferation. Secondly, while our studies provide evidence that loss of CASC15 expression reduces fibrous cap formation and the abundance of lesional ACTA2+ cells, the role of CASC15 in modulating SMC phenotypic fate in atherosclerosis warrants further evaluation. In particular, scRNAseq and scATAC-seq suggest enrichment of CASC15 in SMC-derived fibromyocytes. Our present studies should be complemented with SMC fate mapping and disease stage-dependent loss- and gain-of-function to further delineate the role of CASC15 in atherosclerosis pathogenesis.

CASC15 deficiency not only reduces proliferation but also induces SMC hypertrophy, which has direct pathophysiological relevance to hypertension. Thickening of resistance and conductance arteries is a major pathological feature of systemic hypertension, caused by increased SMC mass. Pioneering studies by Gary Owens’ group have established that hypertension-induced aortic remodeling was primarily due to SMC hypertrophy rather than hyperplasia^13,14,63^. SMC hyperplasia has been predominantly linked to constrictive remodeling of resistance arteries. However, electron microscopy revealed that hyperplasia and hypertrophy co-occur in small arteries^15,16^. Our studies linking SMC hypertrophy to ANG-II-mediated CASC15 deficiency suggest that this lncRNA might play a central mechanistic role in hypertension. CASC15 KO mice display baseline conductance and resistance artery hypertrophy and vasoconstriction. This phenotype likely results from a combination of increased smooth muscle cell (SMC) mass and heightened cytoskeletal tension, aligning with established links between hypertrophic remodeling and reduced vascular compliance^90^. Our RNA-seq analysis further supports these findings, revealing profound disruptions in cytoskeletal and actin organization. While ACTA2 and MYH11 are standard contractile markers, their dysregulated upregulation in CASC15-deficient mice appears maladaptive, contributing to impaired architecture rather than physiological function. These data suggest that CASC15 is essential for maintaining SMC functional competence and preventing pathological transitions. Of major interest, SNPs in *CASC15* have been associated with coronary artery disease and blood pressure regulation^42,44,46,91^. Yet, the impact of these SNPs on CASC15 expression remains to be evaluated. SNPs may also bias the expression of disease-specific CASC15 isoforms. Supporting this hypothesis, several *CASC15* SNPs associated with blood pressure are in splicing regions (**Supplementary Table 3**). Together, our studies suggest that CASC15 might be a central regulator of SMC participation in vascular disease.

Mechanistically, previous studies have associated CASC15 with cell- and context-dependent miRNA sponging or chromatin remodeling^53,57^. Our data support a post-transcriptional model in which CASC15 acts through RNA-binding proteins and RNA-processing complexes rather than through inferred distal chromatin occupancy. This distinction is important given recent evidence that probe-based RNA–chromatin occupancy methods, including ChIRP-seq, CHART-seq and RAP-seq, can be confounded by RNA-independent recovery of genomic DNA fragments and may overestimate the prevalence of trans-acting lncRNA–chromatin interactions^92^. Interaction of CASC15 with small nuclear riboproteins (SNRPs) and ribosomal proteins (RPSs and RPLs) was notable. Nucleolin was also identified as a CASC15 interacting protein. Nucleolin, a multifunctional nucleolar RNA-binding phosphoprotein involved in many RNA processes, including splicing, stability, and rRNA assembly. Nucleolin has been shown to associate with components of the spliceosome machinery and modulate alternative splicing by binding to specific intronic and exonic regulatory sequences and recruiting splicing factors^93,94^. It can also promote RNA stability through RNA-binding domains (RBDs) and Arginine-glycine-glycine (RGG) repeat-dependent binding to the transcript 3’UTR, thereby inhibiting its degradation^95^. Finally, Nucleolin can act as an RNA chaperone that facilitates proper pre-rRNA folding and coordinates the ordered recruitment of ribosomal subunits^96^. Nucleolin has previously been associated with SMC phenotypic switching in part through Nucleolin-dependent stabilization of growth factor gene transcripts, including PDGF-BB and EGF^97^. Our studies show that Nucleolin expression is essential for SMC cell-cycle completion, as Nucleolin depletion leads to SMC hypertrophy and polyploidization, phenocopying the effects of CASC15 knockdown in these cells. Importantly, our studies suggest that Nucleolin is essential in mediating CASC15-dependent G2/M checkpoint, as overexpression of CASC15 was insufficient to rescue proliferative capacities in Nucleolin-deficient SMC. Given Nucleolin’s diverse functions, its selective and context-dependent mechanisms of action may be dictated by its interacting partner. In this regard, our modeling studies suggest that CASC15 acts as a scaffold that promotes the coupling of Nucleolin with other RNA-processing proteins, such as SNRPs. This hypothesis is also supported by the fact that Nucleolin interacts with other lncRNAs, including MALAT1 and Fendrr^98,99^. Variation in CASC15 expression levels and/or isoforms might bias the formation of specific RNA processing complexes or select for specific transcripts. Further mechanistic studies are necessary to fully elucidate how CASC15 regulates RNA metabolism. Nevertheless, the CASC15-Nucleolin axis emerges as a critical regulatory node controlling SMC growth fate during vascular remodeling and disease.

## ACKNOWLEDGEMENTS

We thank the Center for Biologic Imaging (supported by NIH 1S10OD019973), at the University of Pittsburgh, the Health Sciences Sequencing Core at Children’s Hospital of Pittsburgh, the Health Sciences Library System at the University of Pittsburgh, and the Flow Cytometry Core. We thank the Wayne State University Proteomics Core, which is supported through NIH grants P30ES036084, P30CA022453, and S10OD030484. We thank Mike Calderon, Jonathan Franks, Mara Sullivan, and Claudette St Croix from the Center for Biologic Imaging for their contributions to vessel imaging and morphometry analysis. We thank Dr. Satoshi Okawa from the University of Pittsburgh for his assistance with the CASC15 SNP dataset. We thank Drs. St. Hilaire, Straub, and Becker laboratories for providing valve interstitial cell, endothelial cell, and cardiomyocyte RNA extracts, respectively. The schematics were created with BioRender.

## AUTHOR CONTRIBUTION

DG, CED, and IAA conceived the project. DG, CED, IAA, and TBN designed the experiments. IAA, LR, SR, JW, SME, AB, AK, SO, and CED performed experiments and analyzed the data. DG and IAA wrote the manuscript. All co-authors edited the final version of the manuscript.

## DATA AVAILABILITY

RNA-seq data generated in this study have been deposited on the NCBI GEO Datasets platform (accession number: GSE336721) and are publicly available as of the date of publication.

## METHODS

### Mice

All animal protocols were reviewed and approved by the University of Pittsburgh Institutional Animal Care and Use Committee. Mouse housing and experimental procedures were conducted at the University of Pittsburgh’s animal facilities, accredited by the American Association for Laboratory Animals. Mice were housed under a 12-hour light/dark cycle, with controlled temperature (20-26°C) and humidity (50%-60%). Animals had unlimited access to water and a standard rodent chow diet. All mice were backcrossed onto a C57BL/6J background. SMC fate-mapping mice (*Myh11*-CreER^T2^-YFP) were obtained by crossing *Myh11*-CreER^T2^ (Jackson Laboratory #019079) and R26R-EYFP (Jackson Laboratory #006148) mice, as previously described^21^. Only male genotyped mice were used for the experiment because the *Myh11*-CreER^T2^ transgene is located on the Y chromosome. Cre-mediated recombination was initiated at six weeks of age with a series of daily tamoxifen injections (10 injections; 100 µL at 10 mg/mL) dissolved in peanut oil over a 2-week period. The CASC15 KO mouse line was generated by the University of Pittsburgh Transgenics and Gene Targeting core. In brief, a CASC15 null allele was generated using SpyCas9, and 2 target sites were identified within the region (chr13:28884410-28885925, mm10 assembly) using CRISPOR: TTTCCCCGGGCGTGAGAGCGCGG and AAAGTATCCTAGCGCCTCGCAGG^100^. A mixture of the Cas9 protein and sgRNAs was microinjected into fertilized embryos (C57BL/6J, The Jackson Laboratory). Identification of potential founders (F0) was performed by PCR via genotyping the crude DNA lysate obtained from the clipped ear. Validation of CASC15 KO is presented in **Supplementary Figure 15**. Male and female mice were used for experiments.

### Cell Culture

Mouse and rat aortic smooth muscle cells (mSMC) were cultured in growth medium (DMEM: F12, Gibco, 11320-033) supplemented with fetal bovine serum (10%, Corning, 35-015-CV), L-glutamine (1.6 mM, Gibco, 25030081), and penicillin-streptomycin (100 U/mL, Gibco, 15140122) in a humidified sterile incubator at 37 C with 5% CO_2_. For all experiments, mouse and rat SMCs were starved in serum-free media supplemented with L-glutamine (1.6 mM, Gibco, 25030081), Apo-Transferrin (5 mg/ml, Sigma Aldrich, T5391), L-ascorbic acid (0.2 mM, Sigma Aldrich, A4403), and Na Selenite (6.25 ng/ml, Sigma-Aldrich, S5261) for 48 hours, as previously described^19^. Human aortic smooth muscle cells (HAoSMCs; Thermo Fisher Scientific, Cat. No. C-007-5C) were cultured according to the manufacturer’s recommendations in Smooth Muscle Cell Growth Medium 2 (PromoCell; components C-22262 and C-39262) supplemented with 10% fetal bovine serum and the appropriate growth supplements. Cells were maintained at 37 °C in a humidified incubator with 5% CO₂ and used between passages 4 and 7.

### Reagents and Preparation

Human Platelet-Derived Growth Factor-BB (PDGF-BB; Sigma Aldrich, SRP3138) recombinant was reconstituted in 10 mM Acetic Acid and used at 10-30 ng/ml for treatment. Human Angiotensin II (Sigma Aldrich, A9525-50MG) recombinant was reconstituted in distilled water and used at 10ng/ml for treatment. Angiotensin 1 Receptor inhibitor (Olmesartan, MCE, RNH-6270) was reconstituted in DMSO and used at 10nM. BrdU, 5-Bromo-2′-Deoxyuridine, (Fisher, B23151) was reconstituted in DMSO at 10 mM. Antisense LNA GapmeR against mouse CASC15 (Qiagen, 339511-LG00859208-DDA and 339511-LG00250400-DDA) and control (Qiagen, 339515-LG00000002-DFA) were reconstituted in ddH_2_O at 10 mM. Antisense LNA GapmeRs against human *CASC15* were designed to target the conserved region (E23-E28). HAoSMCs were transfected with GapmeR B at a final concentration of 50 nM using Lipofectamine in Opti-MEM.

Mouse Nucleolin siRNA duplexes (2 nmol each; OriGene SR419361) and Trilencer-27 Fluorescent-labeled transfection control siRNA duplex (1 nmol; OriGene SR30002) were reconstituted with 100 uL and 50 uL ddH_2_O at 10 mM, respectively.

### Transfection

GapmeR and plasmid transfections were performed using Fugene HD transfection reagent (Promega, PRE2311) and Opti-MEM reduced-serum medium (Fisher, 31985062). siRNA transfection was performed with Lipofectamine RNAiMax transfection reagent (Invitrogen, 31985062) and Opti-MEM reduced serum medium (Fisher, 31985062). Gapmers and plasmids DNA were transfected at 1ug/mL while siRNA control and nucleolin constructs were transfected at 10 ng/mL. All transfections were performed for at least 24 hours prior to the experimental endpoints.

### CASC15 isoform characterization

A total of 267 transcript isoforms of the 2610307P16Rik gene were annotated at the time of the cloning (source Ensembl Genome Browser 115). Mouse aortic SMC cDNA was prepared by reverse transcription from DNase I-treated RNA. Systematic exon PCR amplification was performed, followed by gel separation, gel extraction (NEB: Monarch T1020S), cloning in pcDNA3, and Sanger sequencing. Full-length isoforms were generated following the same pipeline and fully sequenced. Exon and junction-specific PCR was performed using human SMC cDNA pre-treated with DNAse 1. Isoform-specific qPCR was used to determine dominant isoforms expressed in HASMC. The primer sequences used for isoform characterization are included in **Supplementary Table 13**. The 2D and 3D structures of the CASC15 transcripts were predicted using Vienna Suite and AlphaFold3^101^.

### Real-time quantitative PCR (RT-qPCR)

Cultured SMCs were collected and total RNA extraction was carried out by Qiazol Reagent (Qiagen, 79306), genomic DDNA removal with DNase I and RNA extraction mini (Zymo Research) according to the manufacturer’s protocol. For in vivo RNA extraction, flash frozen tissues were crushed and homogenized in Qiazol Reagent (Qiagen, 79306) by sonication. RNA extraction with DNase I treatment was performed with RNeasy Mini Kit (Qiagen, 74104) following the manufacturer’s instructions. RNA concentration was quantified by Qubit RNA Broad Range Assay kit (Invitrogen, Q10210). Subsequently, reverse transcription reaction was performed to generate cDNA from 1000 ng of RNA using iScript cDNA Synthesis Kit (Bio-Rad, 1708891) according to the manufacturers protocol. RT-qPCR was performed with PowerUp SYBR Green Master Mix (Applied Biosystems, A25742) in a CFX Connect Realtime System machine (Bio-Rad, 1855201). mRNA levels of target genes were normalized to 18s or Gapdh expression from the same independent experiments. The primer sequences are listed in **Supplementary Table 13**.

### Hypertrophy Assay

SMCs were seeded at 10,000 cells/coverslip and placed in 6-well plates in growth medium for 24 hours. 24 hours after transfection, the cells were starved for 48 hours, then treated with vehicle or ANGII for 24 hours. Cells were then fixed (2% PFA) and permeabilized with 0.1% Triton X-100. Cells were thrice washed with 0.01% Triton X-100 permeabilization buffer and blocked with horse serum. Alexa-488 or Alexa-647 directly conjugated phalloidin antibody (1/250) and DAPI (1/1000) were used to stain the filamentous actomyosin cytoskeleton and nuclei, respectively. The solution was incubated at room temperature for 1 hour. Stained cells were washed with 0.01% Triton X-100 buffer and 1X PBS before being imaged with fluorescence microscopy (Leica DMi8). Images were analyzed using ImageJ to measure cell area (based on phalloidin staining), nuclear area (based on DAPI staining), and binucleation rate (based on merge).

### Cell cycle & Polyploidy analysis based on Propidium iodide-DAPI incorporation

SMCs were seeded at 95,000 cells per 6-well-plate well in growth media for 24 hours. Cells were transfected with Control- or CASC15-GapmeR for 24 hours, and then serum-starved for 48 hours. SMCs were trypsinized into a single-cell suspension and collected into a cell pellet by centrifugation at 500g. Trypsinized cells were fixed at −20°C with 70% cold ethanol for at least 16 h before staining. DNA content was stained with a Propidium Iodide (PI) Flow Cytometry Kit (Abcam, ab139418) according to the manufacturer’s instructions. Cell cycle phase and ploidy were determined by FACS in a BD LSR II Flow Cytometer at the University of Pittsburgh Unified Flow Cytometry Core. Data was acquired and analyzed on FlowJo.

### In vitro Proliferation Assay

SMCs were seeded at 10,000 cells/coverslip and placed in 6-well plates in growth medium for 24 hours. After cells were transfected for 24 hours, mSMCs were serum-starved for 48 hours. After this, the cells were treated with acetic acid (control) or PDGF-BB and 10uM BrdU for 24h before being harvested for analysis. Proliferation was assessed by BrdU incorporation quantification. Briefly, after incubation with BrdU (24 hours), mSMCs were washed 3 times with PBS and fixed with 2% PFA/PBS for 15 min. Washed and fixed cells were permeabilized with 0.1% Triton X-100 for 20 minutes at room temperature. The cells were treated with the sequential addition of cold 1N HCl and room-temperature 2N HCl for a total of 30 mins. Phosphate/citric acid buffer, pH 7.4, was added and incubated for 10 minutes. Cells were thrice washed with 0.01% Triton X-100 permeabilization buffer and blocked with antibody staining buffer (PBS in FBS/Goat or horse serum). Anti-rat BrdU primary antibody (Abcam, ab6326) at 1/250 was added overnight. After washing with 0.01% Triton X-100 permeabilization, Alexa Flour donkey anti-rat secondary antibody (Invitrogen A-21208, 1:250) and DAPI (1:1000) were added. The antibody mix was incubated for one hour at room temperature. Stained cells were washed with 0.01% Triton X-100 permeabilization buffer and 1X PBS before being imaged with a fluorescence microscopy.

### Scratch assay (Migration)

SMCs were seeded at 9.5*10^4^ cells per well to achieve 60-70% confluency in growth medium into 6-well plates for 24 hours. After cells were transfected for 24 hours, mSMCs were serum-starved for 48 hours. A 2-mm-wide comb was used to make a uniform scratch or wound at the center of the culture plate. Cells were then treated with acetic acid (control) or PDGF-BB for 24 hours. Images were collected at 0, 8 and 24 h on a fluorescent microscope (Leica, DMi8) using the Ocular Advanced Scientific Camera Control software (Digital Optics Limited). ImageJ was used for Image processing and wound closure measurement.

### Histology

#### Electron microscopy

SMCs were seeded at 2 x 10^5^ cells per 35 mm × 10 mm dish to achieve an 80-90% confluent monolayer in growth medium after 48 hours. The cells were then transfected for 24 hours with Control or S-CASC15 GapmeRs. 24 h post-transfection, SMCs were serum-starved for 48 h and washed 3 times with 1x PBS. The cells were fixed in situ with 5 mL cold 2.5% glutaraldehyde in 1x PBS, pH 7.4. The dishes were transferred to the University of Pittsburgh Center for Biological Imaging for transmission electromagnetic processing and imaging. Briefly, the specimens were rinsed in PBS, post-fixed for 1 hour at 4 °C in 1% OsO_4_ with 1% potassium ferricyanide, and then washed thrice with 1x PBS. Dehydration procedures were performed through a graded series of ethanol and embedded in Poly/Bed® 812 (Glauert formulations- Dodecenyl Succinic Anhydride, Nadic Methyl Anhydride, Poly/Bed 812 Resin, and Benzyl dimethyl amine). Ultrathin sections (65 nm) were cut on a Leica Reichert Ultracut (Leica Microsystems, Buffalo Grove, IL) and stained with Uranyless and Reynold’s Lead Citrate. The specimens were examined on a JEOL 1400-PLUS 120kV transmission electron microscope, and images were acquired with a side-mount Nikon 50M digital camera (Nikon, Melville, NY).

#### Immunocytochemistry

For in vitro immunofluorescent staining, cultured cells were fixed with 4% PFA/PBS (Electron Microscopy Sciences) and permeabilized with 0.2% Triton X-100 (Sigma). Staining was performed for actin fiber using Alexa Fluor Phalloidin-647 or −488 (Invitrogen A22287, A12379) and DAPI (1:1000).

#### Immunofluorescent staining

All collected tissues were vertically embedded. After that, serial 10mm sections were collected. For ligated tissues, sections covering 180mm from the ligation site, starting at the level of the visible suture, were used for immunofluorescent staining of tissue slides. Obtained sections were deparaffinized, and antigen retrieval was performed by the heat method (Vector Laboratories H-3300). Tissue staining was performed using primary antibodies for GFP (Abcam ab6673, 1:250) or control IgG from same species, and Alexa Fluor donkey anti-goat 555 (Invitrogen, 1:250) as secondary antibody; ACTA2 (ab5694, 1:250) and Alexa Fluor donkey anti-rabbit 555 or 647 (Invitrogen A31572, 1:250); anti-goat CD31 (R&D AF3628, 1:100). Where applicable, conjugated antibodies for ACTA2-FITC (Sigma Aldrich, F3777), WGA-640 (Biotum 29026-1, 1:2000) were used. Along with the secondary antibodies, sections were also stained for ACTA2 (Sigma-Aldrich, F3777, 1:500) and DAPI (1:1000) or Hoechst (Invitrogen, 1:250). Slides were mounted using Prolong Gold Antifade Reagent (Fisher P36930). For in vivo proliferation studies with EdU, multiplexing with antibodies was performed with Click-iT™ Plus Alexa Fluor™ 647 Picolyl Azide Toolkit (ThermoFisher, C10643) according to the manufacturer’s instructions.

#### Image acquisition

Images were acquired on a fluorescence microscope (Leica DMi8) using the Ocular Advanced Scientific Camera Control software (Digital Optics Limited) or on a Nikon A1 Confocal microscope using NIS-Elements software (Nikon). Image processing was performed using ImageJ.

### RNA sequencing and data analysis

Total RNA was extracted and column-purified using the RNeasy Mini Kit (Qiagen, 74104) based on the manufacturer’s protocol. Genomic DNA was removed using DNase I (Qiagen). RNA quantification was performed using Nanodrop (ThermoFisher). RNA quality control, library preparation, and sequencing was performed by NOVOGENE. Briefly, RNA quality was assessed with the Agilent RNA Screen Tape Assay Tape Station System. RNA library was prepared by TruSeq Stranded mRNA (PolyA+) according to the manufacturer’s protocol. Libraries were sequenced on an Illumina NextSeq High Output 150-cycle kit (2 x 75 bp), yielding a depth of 20 million reads per sample. Preprocessing was conducted with nf-core/rnaseq: v3.22.2’s standard parameters unless otherwise stated. The 75 bp paired-end reads were mapped to Mus musculus reference genome, GRCm39 primary assembly with the STAR aligner v2.7.11b and quantified with RSEM 1.3.1. Differential Gene Expression Analysis (DGEA) was conducted on the gene count matrices in R using DESeq2; genes with a p-adjusted value < 0.05 and a fold change > 2 were considered differentially expressed genes (DEGs) and plotted with the EnhancedVolcano package. Gene Ontology (GO) analysis was conducted on DEGs with clusterProfiler’s enrichGO function using all aligned genes as the universe with pvalueCutoff = 0.05 and qvalueCutoff = 0.10. GO analysis was plotted in GraphPad. Gene Set Enrichment Analysis (GSEA) was conducted on DEGs and plotted with enrichplot’s gseaplot function. The pheatmap library was used to plot heatmaps of counts data, either of transcript per million (TPM) or fold change (FC) vs. controls. LncRNA data analysis was performed on previously published RNAseq data (NCBI GEO Accession number: GSE179220) (Liu et al., 2021). The lncRNA analysis and data plotting methods are the same as those stated for RNA-seq. Cut-offs are fold-change > 1.7 and p value < 0.05.

### Right Carotid artery Ligation (RCL)

Unilateral ligation of the right carotid artery in *Myh11*-CreER^T2^ YFP mice was performed as detailed in ^19,23^. At 6 weeks of age, male mice were injected daily with 1mg of tamoxifen in peanut oil (10 mg/mL) for 10 days. 7 to 14 days after the last tamoxifen injection, mice were anesthetized with isoflurane, and the right carotid artery was ligated caudal to the bifurcation using 7-0 silk suture. At the time of the ligation, pluronic gel (20% Wt/Vol) containing lentiviral particles (10^6^-10^7^) encoding ShRNA-Control, ShRNA-CASC15, pLVX, or pLVX-CASC15 was applied on the injured artery. The lentiviral particles were unilaterally delivered and applied to the ligation site using Pluronic gel. After 28 days, all experimental mice underwent euthanasia by CO_2_ asphyxiation and were perfused with 4% PFA/PBS (4% paraformaldehyde, Electron Microscopy Sciences in 1X phosphate buffered saline, Gibco) with a gravity perfusion system. Left and right carotid arteries were excised, placed in 4% PFA/PBS overnight, processed, and vertically embedded in paraffin. 10 mm tissue sections were collected from the bifurcation suture onto staining slides using a microtome.

### Atherosclerosis studies

Atherosclerosis was induced using an adeno-associated virus-mediated PCSK9 overexpression model. CASC15^+/+^ and CASC15^-/-^ mice were administered a single tail vein injection of AAV8-D377Y-mPCSK9 (Vector Biolabs) at a dose of 1.75 × 10^12 genome copies (GC) per mouse. Following a 2-day recovery period, mice were fed a Western diet (TD.88137, Envigo; 42% kcal from fat) for 16 weeks. At the study endpoint, mice were euthanized, perfused with 4% PFA, and the brachiocephalic artery (BCA) was carefully dissected and cleared of surrounding adipose and connective tissues. BCAs were embedded in paraffin, and BCA 10-µm-sections were collected from the junction with the aortic arch^102^. Atherosclerotic lesion formation was evaluated by histological analysis of arterial cross-sections.

### Ex-vivo myography

Following euthanasia and systemic perfusion with PBS, thoracic aortas and mesenteric resistance arteries were harvested from CASC15^−/−^ and CASC15^+/+^ mice, carefully cleared of surrounding adipose and connective tissue, and sectioned into 2-mm rings. Aortic rings were mounted on pins, whereas mesenteric artery rings were mounted on two 40-μm stainless-steel wires in a myograph system (DMT 620M, Danish Myo Technology) for isometric tension recordings. Vessels were maintained in physiological salt solution (PSS; NaCl 119 mM, KCl 4.7 mM, MgSO_4_ 1.17 mM, KH_2_PO_4_ 1.18 mM, D-glucose 5.5 mM, NaHCO_3_ 25 mM, EDTA 0.027 mM, and CaCl_2_ 2.5 mM) at 37°C and continuously aerated with 95% O_2_ and 5% CO_2_. After a 30-min equilibration period, mesenteric arteries were normalized according to the manufacturer’s guidelines, whereas aortic rings were equilibrated at a resting tension of 5 mN prior to functional studies. The vessels were constricted by the addition of 60 mM potassium solution (KPSS) for 5 minutes, followed by concentration-response curves to phenylephrine (10^-10^–10^-4^ M) and U46619 (10^-10^–10^-5^ M).

### SMC morphometric image analysis

Morphometric analysis for in vitro studies was carried out using 10 fields of view per technical replicate. Analysis of cell area, nuclear area, nuclear counts, and BrdU index was performed using ImageJ. Binucleation was estimated as the number of nuclei per cell. BrdU index ratio or percentage was estimated as the ratio of BrdU-positive cells to the total number of nuclei. For in vivo analysis, vessel morphometry was performed by designing nodes to identify SMC-specific nuclei using the DAPI channel and ACTA2 expression, while excluding nuclei colocalizing with CD31 staining. Cell area was determined based on WGA staining. Morphometric parameters such as vessel width, media width, SMC area (WGA), and nuclear area (DAPI) were measured using automatic detection based on adequate staining.

### RNA fluorescence in situ hybridization (RNA-FISH)

RNA-FISH was performed on cultured mSMCs using the miRCURY LNA miRNA ISH Detection system (Qiagen) according to the manufacturer’s instructions with minor modifications. SMCs were seeded onto glass coverslips and transfected with either an empty vector (pcDNA) or a CASC15 overexpression plasmid (pcDNA-CASC15). Forty-eight hours after transfection, cells were washed with phosphate-buffered saline (PBS), fixed in 4% paraformaldehyde for 15 min at room temperature, and permeabilized with 0.1% Triton X-100 for 10 min. Hybridization was performed using a custom DIG-labeled miRCURY LNA probe targeting mouse CASC15 (Qiagen, probe ID: LCD0176858-BKG) at a final concentration of 40 nM in hybridization buffer. A scrambled negative control probe and U6 snRNA positive control probe included in the miRCURY LNA miRNA ISH Buffer and Controls kit (Qiagen, Cat. No. 339459) were used to assess probe specificity and assay performance. Coverslips were incubated with probes in a humidified chamber at 65°C under RNase-free conditions. Following hybridization, samples were subjected to a series of stringent SSC washes according to the manufacturer’s protocol. Hybridized probes were detected using anti-DIG-POD antibody (1:400 dilution), followed by signal amplification using TSA Plus Cyanine 3 substrate (PerkinElmer). Nuclei were counterstained with DAPI.

### Nuclear and cytoplasmic RNA fractionation

Nuclear and cytoplasmic RNA fractions were isolated from mSMCs using the Cytoplasmic and Nuclear RNA Purification Kit (Norgen Biotek) according to the manufacturer’s instructions. Briefly, cells were washed with ice-cold PBS and lysed under conditions that preserve nuclear integrity. The lysate was centrifuged to separate the cytoplasmic supernatant from the nuclear pellet. Cytoplasmic RNA was purified from the supernatant, while the nuclear pellet was subsequently lysed to isolate nuclear RNA. RNA from each fraction was purified using the kit columns, including on-column DNase treatment when indicated, and eluted in RNase-free water. RNA concentration and quality were assessed prior to reverse transcription. Equal amounts of nuclear and cytoplasmic RNA were reverse-transcribed and analyzed by quantitative PCR. CASC15 enrichment in each fraction was assessed relative to compartment-specific control transcripts. U6 snRNA was used as a nuclear-enriched control, whereas Gapdh mRNA was used as a cytoplasmic-enriched control. Fraction purity was confirmed by the expected enrichment of U6 in the nuclear fraction and Gapdh in the cytoplasmic fraction.

### LncRNA CASC15 Pulldown

PmeI-linearized biotinylated-CASC15 and biotinylated-antisense-CASC15 transcripts were generated by HiScribe T7 High Yield RNA Synthesis Kit (BioLabs E2040S) according to the manufacturer’s instructions. 1 µg of biotinylated lncRNA CASC15 was denatured by heating to 65 - 80°C for 10 min and cooled down slowly to 4°C^103^. The denatured biotinylated transcripts were added to a low-binding Eppendorf containing 1mg/mL of whole-cell protein lysate. The solution was incubated for at least 12 hours at 4°C on a gentle rotatory shaker. In another low-binding Eppendorf tube, 100 µL of Dynabeads MyOne streptavidin (Invitrogen 65001) was twice washed with ice-cold 1X Binding buffer ( 5 mM Tris-HCl pH 7.5, 0.5 mM EDTA, 1 M NaCl) supplemented with 0.1% (v/v) Tween 20 (Sigma P1379-100ML), complete protease inhibitor cocktail (Sigma 11836170001) and 100 Unit/mL Ribolock RNase inhibitor (Thermofisher EO0381). Samples were twice washed in Ion Solution A (RNase-Free 0.1 M NaOH, RNase-Free 0.05 M NaCl, RNase I), and once in Ion Solution B (RNase-Free 0.1 M NaCl, RNase I). The beads were stored in 2X Binding Buffer with 0.1% (v/v) Tween 20 to a final concentration of 5 ug/uL. The biotinylated probe-bound proteins were isolated using a Dyna-mag magnet for 2-3 minutes, washed 3 times with 1X Washing Buffer + 0.01% Tween, and transferred to a clean 1.5 mL Eppendorf tube before the final wash with 1X Washing Buffer. The solution was digested with mass spectrometry-grade trypsin.

### Mass spectrometry and target identification

Dried peptide samples were reconstituted in 0.1% formic acid and injected for LC–MS/MS analysis. Peptides were separated on a Thermo Scientific Vanquish Neo UHPLC system equipped with an Acclaim PepMap 100 trap column (100 μm × 2 cm, C18, 5 μm, 100 Å) and an Ionopticks Aurora® Ultimate™ XT C18 UHPLC column (75 μm × 25 cm). Chromatographic separation was performed using a 90-minute gradient from 5% to 42% solvent B (80% acetonitrile, 0.1% formic acid) at a flow rate of 270 nl/min, with solvent A consisting of 0.1% formic acid in water. Data-independent acquisition (DIA) was performed on a Thermo Scientific Orbitrap Eclipse mass spectrometer with a nanoelectrospray ionization source operating at +2300 V. MS1 spectra were acquired in the Orbitrap at a resolution of 120,000 over the 350–1400 m/z range, with a normalized AGC target of 750% and a maximum injection time of 50 ms. Peptide fragmentation was induced by higher-energy collisional dissociation (HCD) at 32% collision energy with isolation windows of 16 Da (360–848 m/z) and 32 Da (848–1400 m/z). MS2 spectra were collected in the Orbitrap at a resolution of 15,000 over 200–1600 m/z, with an AGC target of 6000% and a maximum injection time of 50 ms. Raw DIA data were processed using both Spectronaut (v20.1, Biognosys) and FragPipe (v23.1). Spectronaut analysis was performed using the BGS Factory Settings with the UniProt Mus musculus FASTA database (downloaded March 2021; 20,381 entries). Differential expression was determined using an adjusted p-value cutoff of <0.05 and |log2 fold change| ≥ 0.58. FragPipe analysis was conducted with the default setting of DIA_SpecLib_Quant workflow against a reviewed UniProt Mus musculus FASTA database containing decoys and contaminants (downloaded July 2025; 43,842 entries, including 50% decoys). Protein quantification outputs were further analyzed in R (v4.5.0), where principal component analysis was carried out using the prcomp function, and hierarchical clustering with heatmap visualization was generated using the pheatmap package.

### Statistical analysis

All data are presented as the mean ± SEM. In vitro experiments were independently repeated at least 3 times, with duplicate technical replicates. One single data symbol represented the mean value of technical repeats for one independent experiment. All statistics were performed using GraphPad Prism 8. Data were tested for normality using the Shapiro-Wilk test. Two-tailed unpaired Student’s t-tests with a confidence level of 95% were used to compare two groups with continuous variables with normal distributions and equal variances. Two-tailed unpaired Student’s t-tests followed by a Welch’s correction with a confidence level of 95% were performed if the two groups had unequal variances. Two-tailed unpaired Mann-Whitney U-tests with a confidence level of 95% were used if variables were non-normally distributed. For comparison between multiple groups with a single factor or two factors, we used One-way or Two-way ANOVA, respectively. For categorical data, we used two-sided Fisher’s exact test. p < 0.05 was considered statistically significant.

