## Supplementary Figures for "CASC15 dictates vascular smooth muscle cell growth fate and pathological vascular remodeling through post-transcription regulation of mitotic fidelity"

### SUPPLEMENTARY FIGURE LEGENDS

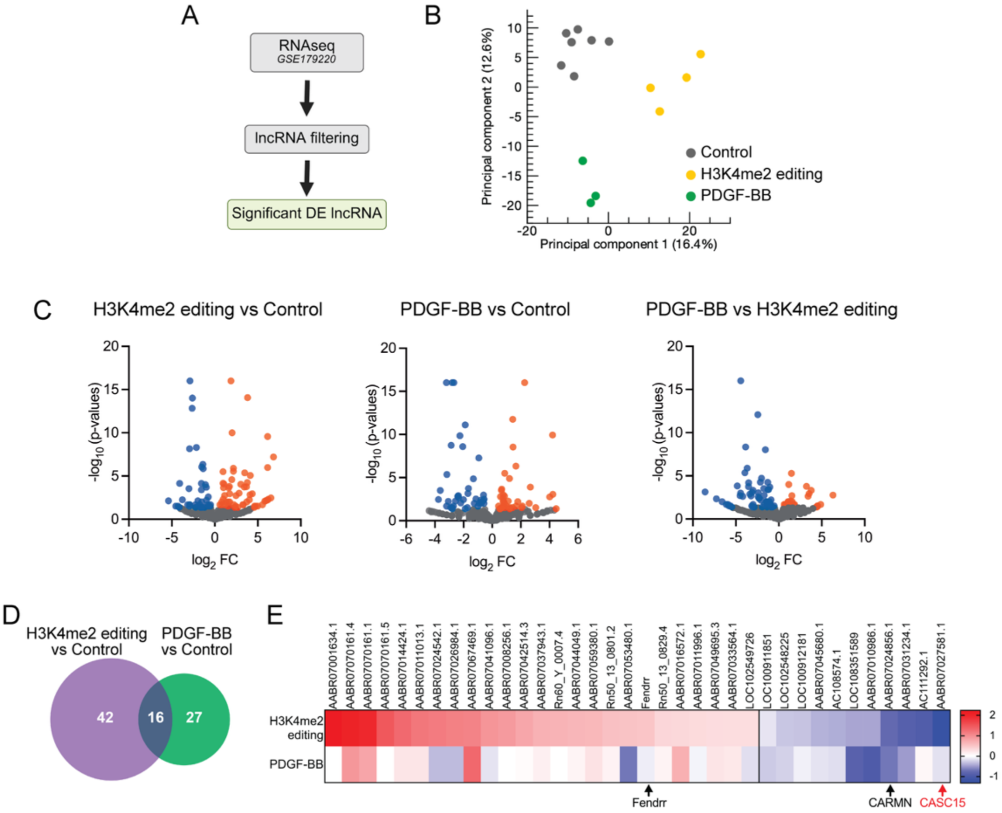


**Supplementary Figure 1: Differential expression of lncRNAs in H3K4me2-edited and PDGF-BB-treated SMCs. A.** Analytical pipeline schematics. **B.** Principal component analysis of bulk RNAseq data obtained on H3K4me2-edited, PDGF-BB-treated, and control SMC (GEO: GSE179220). **C.** Volcano plots representing lncRNA fold change and p-value between groups. **D.** Venn Diagram representing the intersection of differentially expressed lncRNA in H3K4me2-edited and PDGF-BB-treated SMC as compared to control SMC. **E.** Heat map representing lncRNA fold change expression.

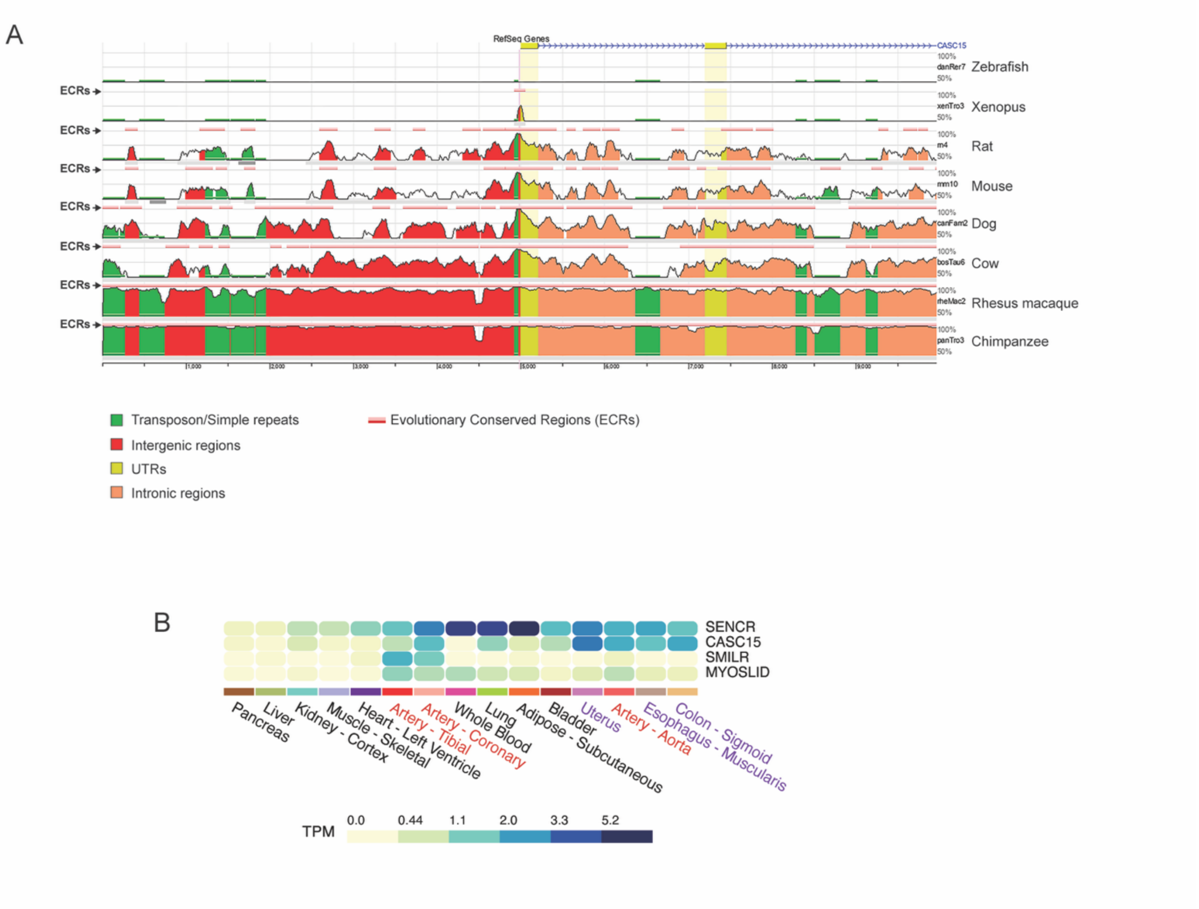


**Supplementary Figure 2: Conservation and SMC-selective expression of CASC15. A.** Genomic conservation of *CASC15* sequence across species. **B.** Expression of CASC15 and other SMC-enriched lncRNA (SENCR, SMILR, MYOSLID) across human tissues (GTEx).

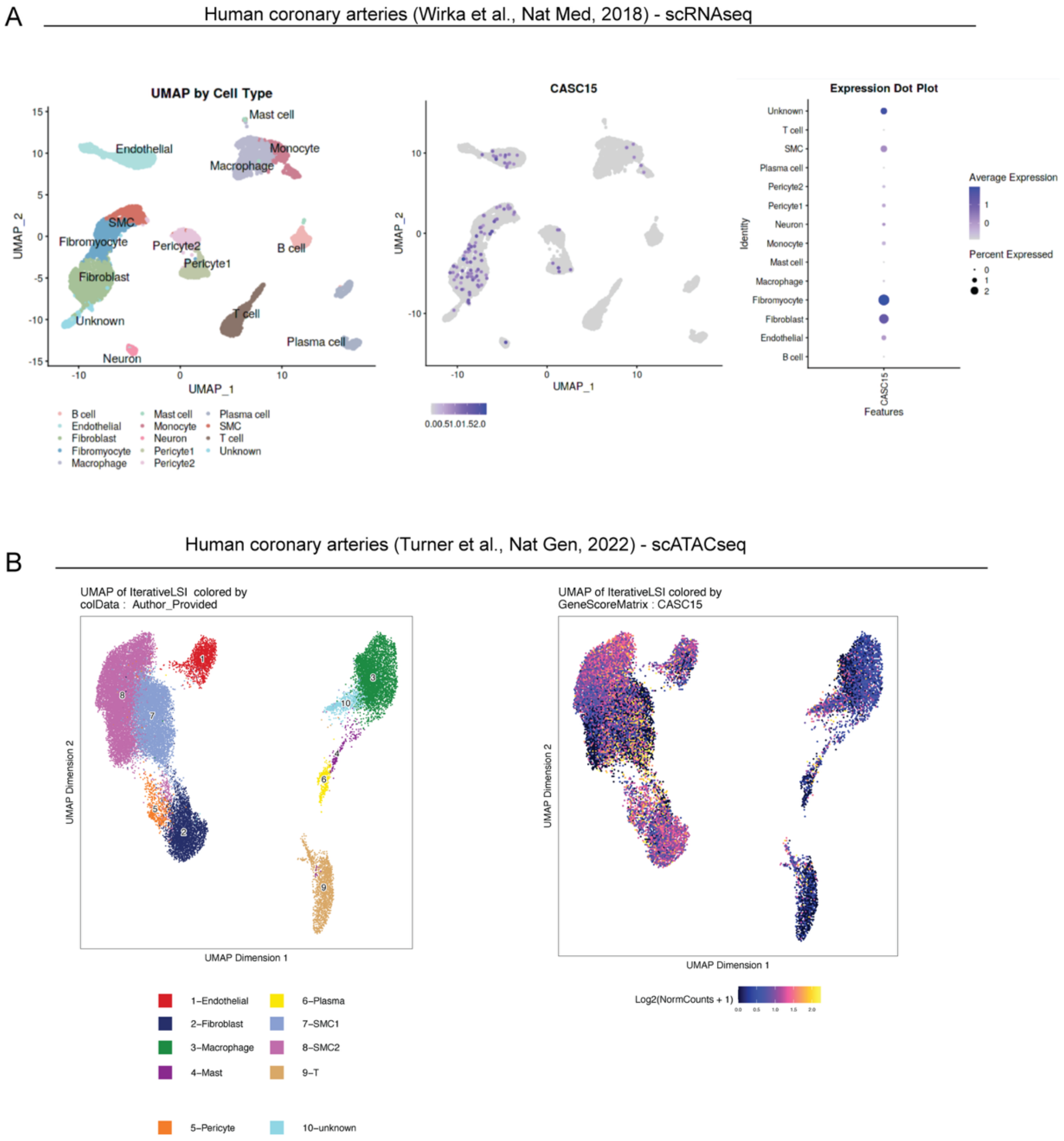


**Supplementary Figure 3: CASC15 expression and chromatin accessibility in human coronary artery disease. A.** Uniform manifold approximation and projection (UMAP) and clustering (left) of human coronary artery cell populations based on single-cell RNAseq (GEO: GSE131780). UMAP plot (center) representing CASC15 expression. Dot plot (right) representing CASC15 average expression. B. UMAP and cell clustering based on single-nucleus ATACseq (GEO: GSE175621) on human coronary arteries. UMAP plots for *CASC15* chromatin accessibility.

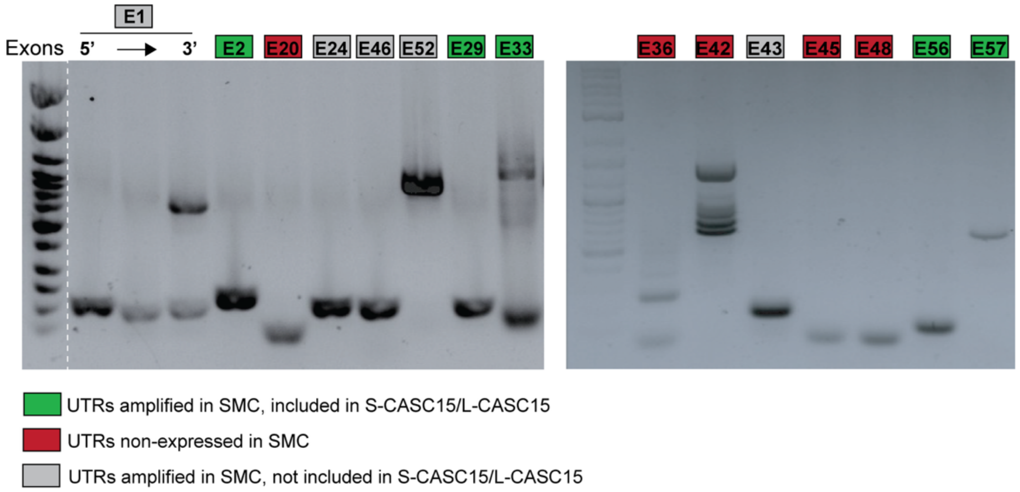


**Supplementary Figure 4: Characterization of 2610307P16Rik (CASC15) untranslated exon expression in mSMC.** Representative gels showing systemic evaluation of untranslated exon expression in RNA extracted from mSMC. While some untranslated regions (UTRs) were not transcribed in mSMC (red), amplified UTRs were either included in S-CASC15 and L-CASC15 (green) or absent from the full-length sequences of these transcripts (gray).

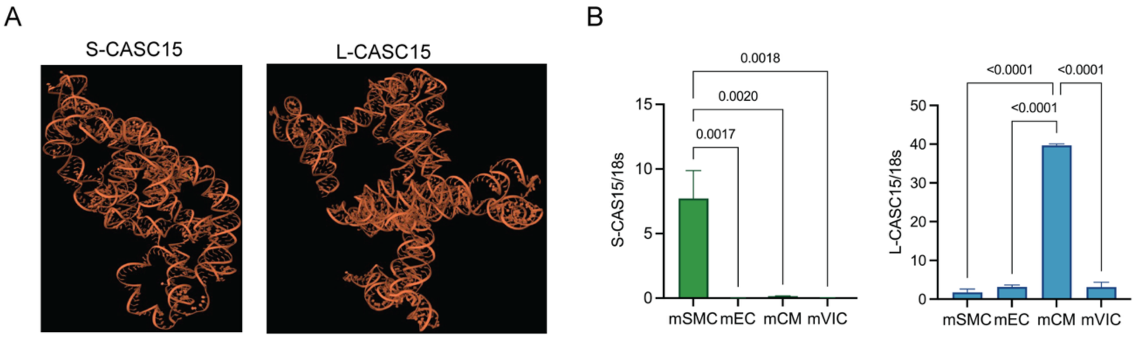


**Supplementary Figure 5: S-CASC15 and L-CASC15 exhibit distinct conformations and lineage-selective expression. A.** S-CASC15 and L-CASC15 conformation prediction using AlphaFold3. **B.** Isoform-specific qPCR quantification of CASC15 isoform expression in mouse SMC, endothelial cells (mEC), cardiomyocytes (mCM), and valve interstitial cells (mVIC). CASC15 expression was normalized to 18s. One-way-ANOVA.

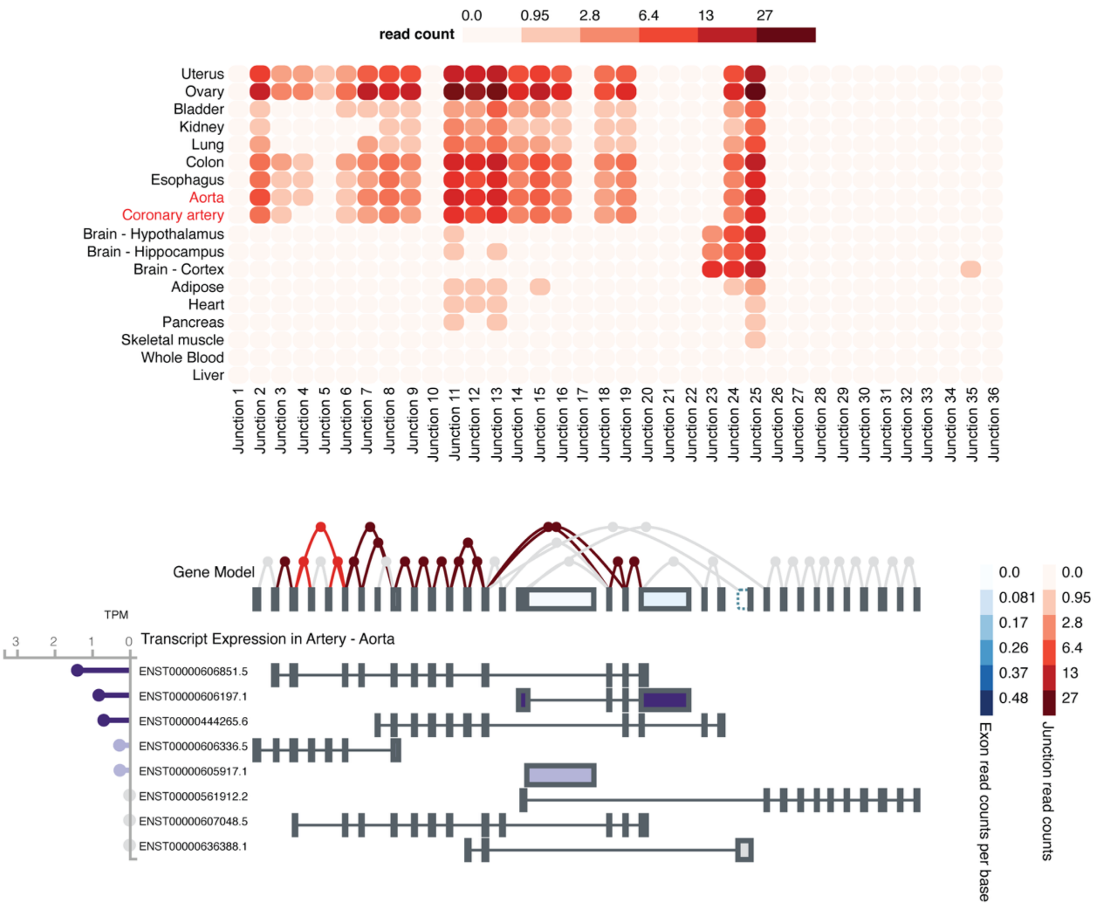


**Supplementary Figure 6: Tissue-specific human CASC15 isoform expression.** Untranslated exon junction read counts in various human tissues (top) and identified transcript variants in the human aorta (bottom). GTEx database.

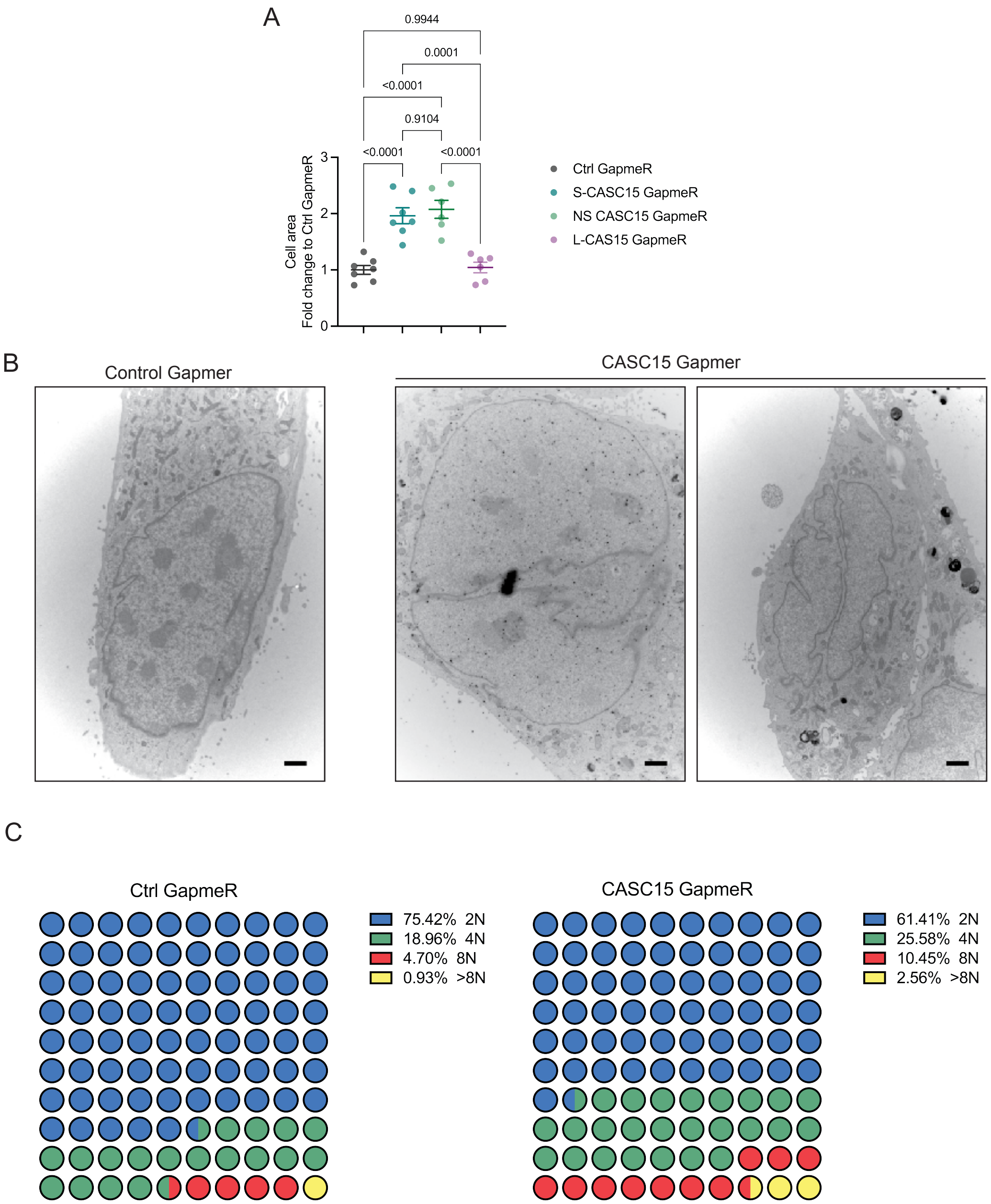


**Supplementary Figure 7: S-CASC15 KD triggers cell hypertrophy, binucleation, and polyploidization. A.** Cell area measurement in SMC transfected with Control, S-CASC15, non-selective (NS) CASC15, and L-CASC15 targeting GapmeRs. Data expressed as fold change compared to Ctrl GapmeR. Data expressed as mean ± SEM. One-way ANOVA. **B.** Representative electron microscopy micrographs of Control and CASC15 GapmeR SMCs showing examples of binucleated cells. Scale bar = 1 µm). **C.** Evaluation SMC ploidy by flow cytometry.

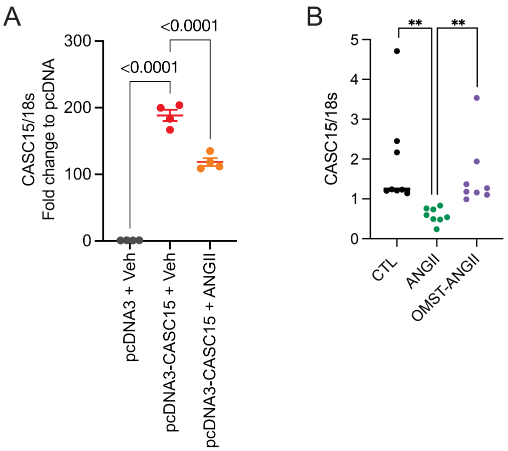


**Supplementary Figure 8: CASC15 expression is negatively regulated by angiotensin II in an AT1-dependent manner. A.** CASC15 expression after transduction with pcDNA-CASC15 and treatment with vehicle or ANGII (100nM, 24h). Data expressed as mean ± SEM. One-way ANOVA. **B.** CASC15 expression in SMC treated with ANGII in absence or presence of Olmesartan (OMST), an AT1 inhibitor. Data expressed as mean ± SEM. One-way ANOVA.

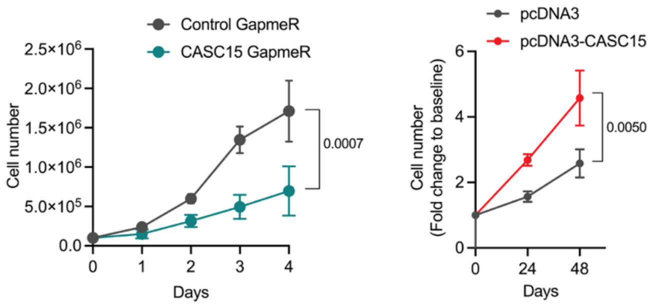


**Supplementary Figure 9: CASC15 regulates cell proliferation.** Cell count in CASC15-deficient (left) and CASC15-overexpressing (right) mSMCs as compared to their respective controls. Data expressed as mean ± SEM. Two-way ANOVA.

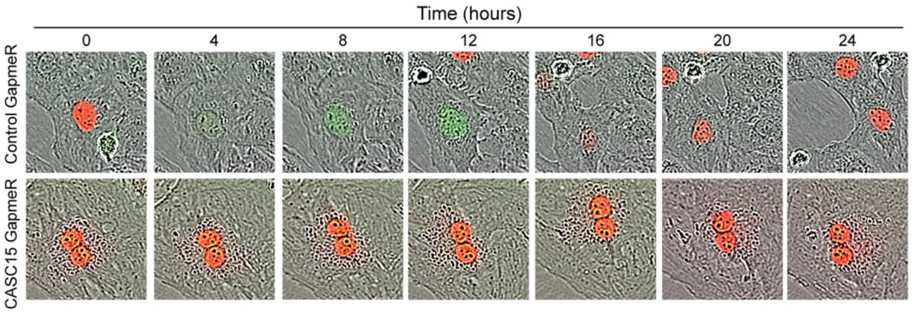


**Supplementary Figure 10: Lack of CASC15 induces polyploidization and G1 arrest.** Live imaging of rat stable FUCCI rat SMCs transfected with Control and CASC15 GapmeR.

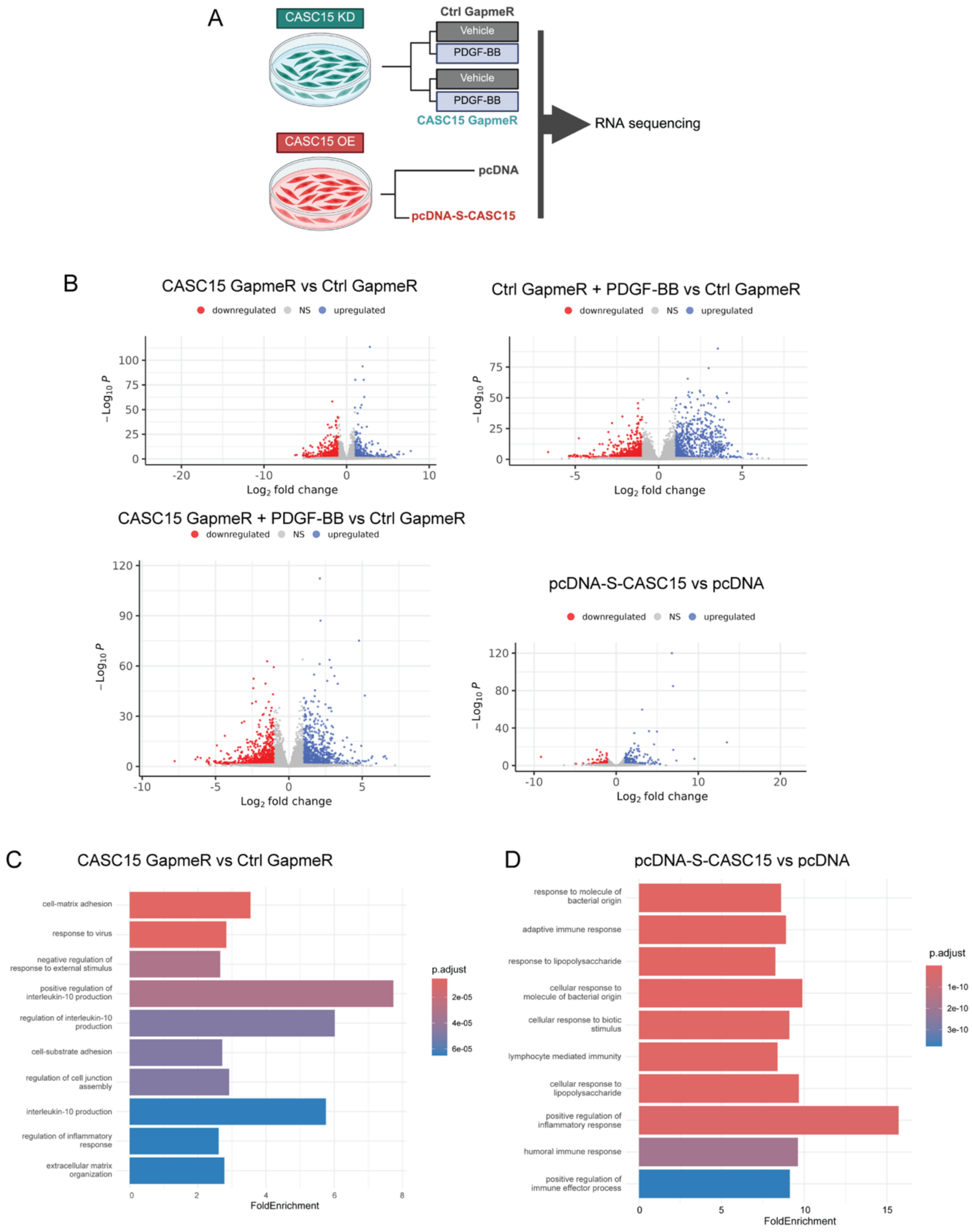


**Supplementary Figure 11: Characterization of transcriptional changes induced by CASC15 loss- and gain-of-function. A.** Schematics representing the experimental groups used for bulk RNA sequencing. **B.** Volcano plots representing differentially expressed genes in experimental groups vs controls. **C.** Top 10 differentially regulated pathways in CASC15-deficient SMC vs control. **D.** Top 10 differentially regulated pathways in CASC15-overexpressing SMC vs control.

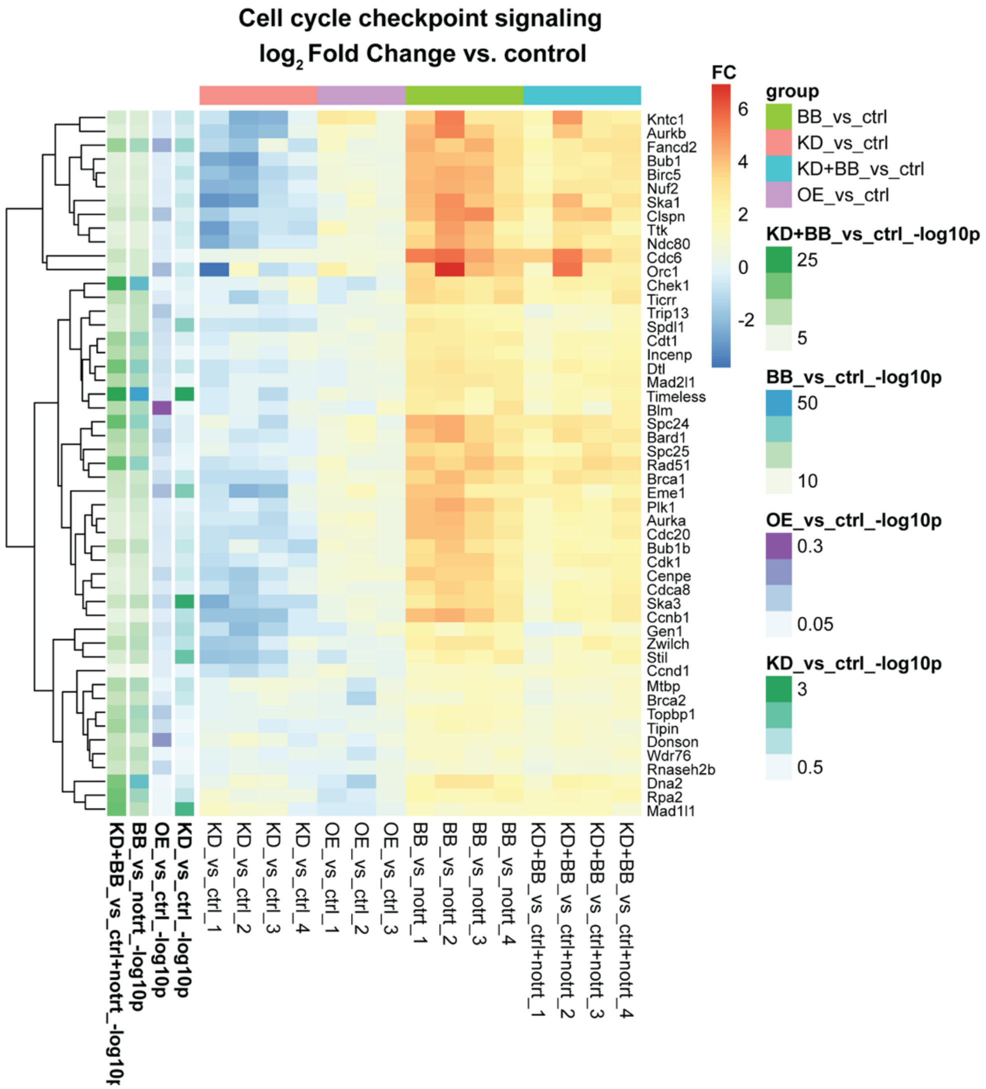


**Supplementary Figure 12: Heat map of genes related to cell cycle checkpoint signaling in experimental groups vs controls.** FC: Fold change.

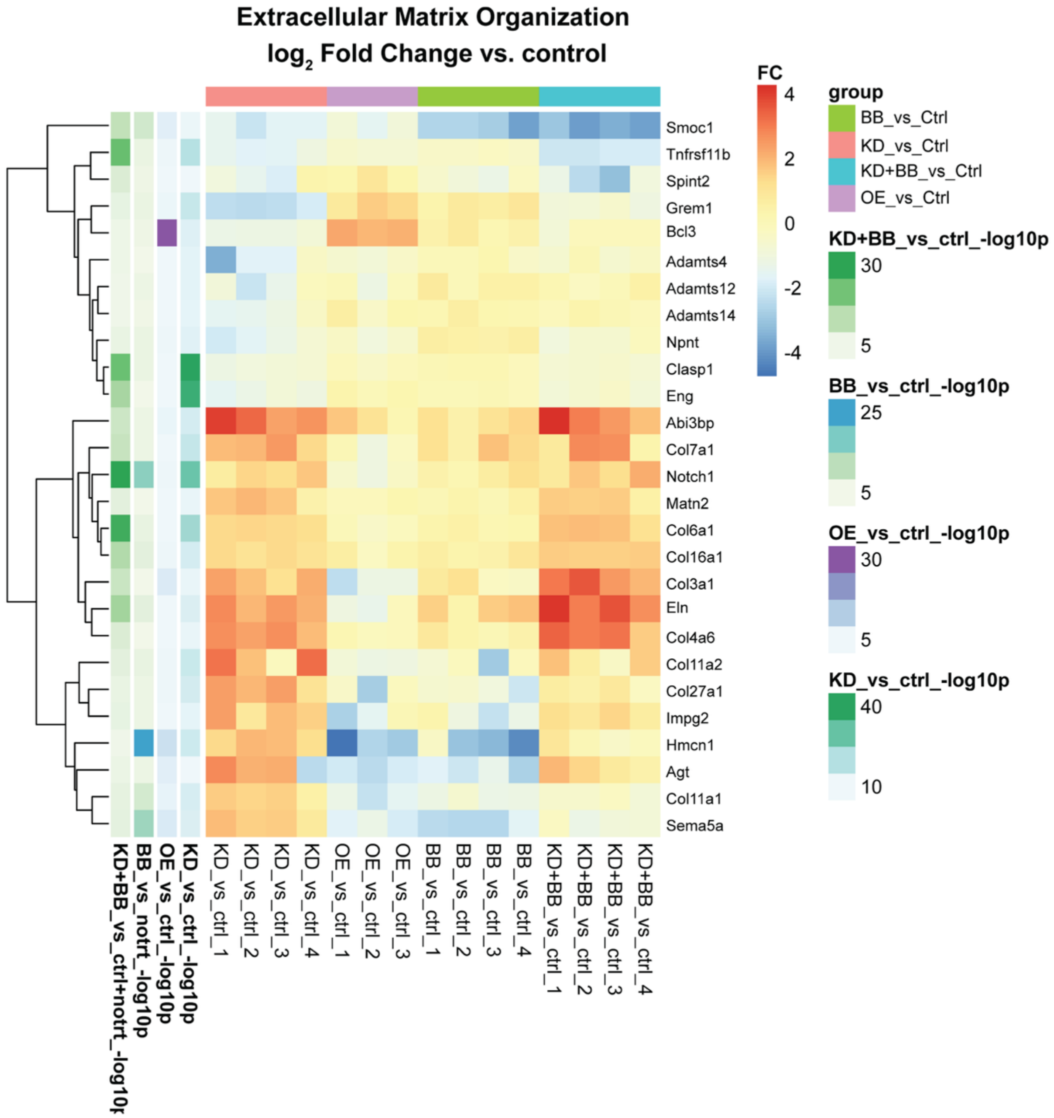


**Supplementary Figure 13: Heat map of genes related to extracellular matrix organization in experimental groups vs controls.** FC: Fold change.

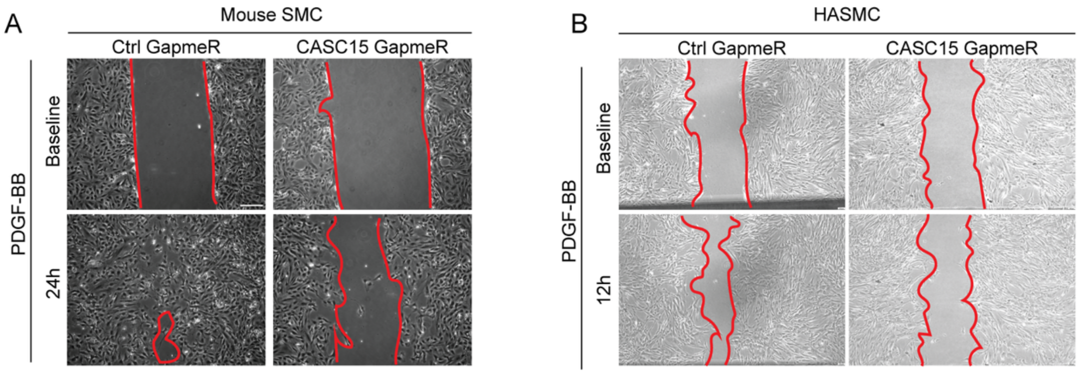


**Supplementary Figure 14: CASC15 KD impairs SMC migration. A.** Representative micrographs of scratch assays using mouse SMC transfected with Control- or CASC15-GapmeR and treated with PDGF-BB (10ng/mL) for 24h. Scale bar = 100 µm. **B.** Representative micrographs of scratch assays using human aortic SMC transfected with Control- or CASC15-GapmeR and treated with PDGF-BB (10ng/mL) for 12h. Scale bar = 100 µm.

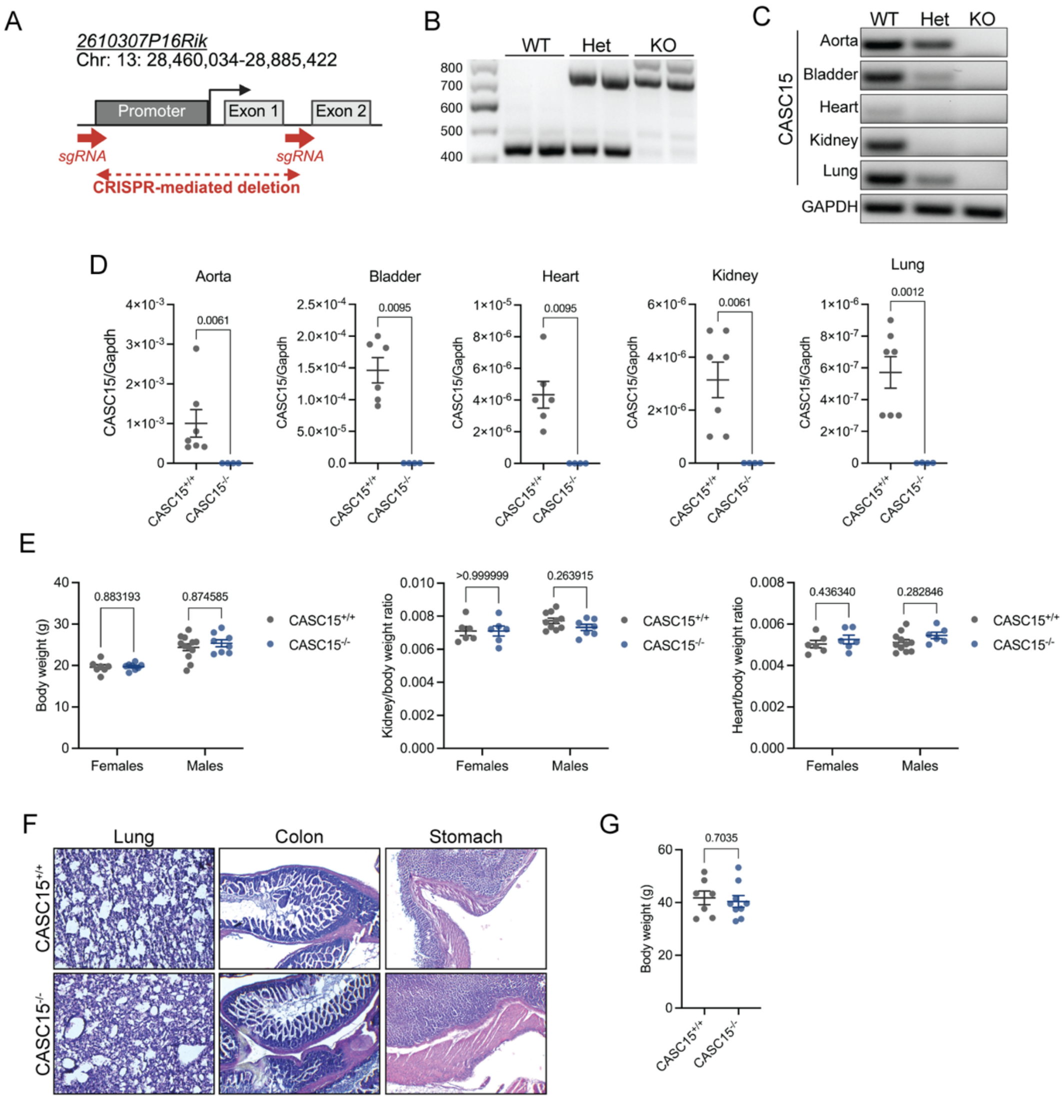


**Supplementary Figure 15: Generation and characterization of a novel CASC15 KO mouse. A.** Schematic representation of CRISPR-Cas9-mediated excision of *2610307P16Rik* proximal promoter, untranslated first exon, and first intron. **B.** Genotyping of CASC15$^{+/+}$, CASC15$^{+/-}$, and CASC15$^{-/-}$ mice. **C.** CASC15 expression (PCR) in CASC15$^{+/+}$, CASC15$^{+/-}$, and CASC15$^{-/-}$ mouse tissues. **D.** Quantification of CASC15 expression by qPCR. Normalized to GAPDH in CASC15$^{+/+}$ and CASC15$^{-/-}$ mice. Data expressed as mean ± SEM.  Student t-test. **E.** Body weight, kidney/body weight, and heart to body weight ratios. Data expressed as mean ± SEM.  Multiple t-test. **F.** Representative micrographs of H&E staining of the lung, colon, and stomach. **G.** Body weight in CASC15$^{+/+}$ and CASC15$^{-/-}$ mice after injection with PCSK9-AAV and 16 weeks of high-fat diet feeding. Data expressed as mean ± SEM.  Student t-test.

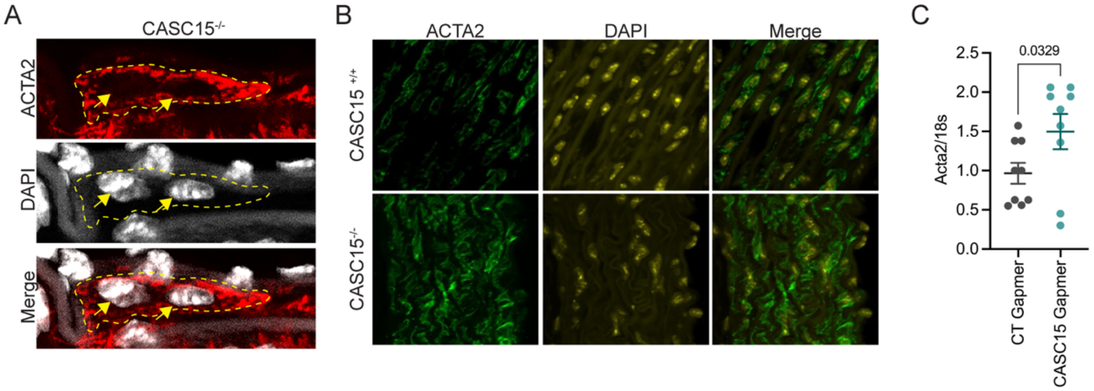


**Supplementary Figure 16: Increased binucleation and ACTA2 expression in CASC15 KO aorta. A.** Example of binucleated SMC in the aortic media of CASC15$^{-/-}$ mice. **B.** Immunofluorescent staining for ACTA2 and DAPI in CASC15$^{-/-}$ and CASC15$^{+/+}$ aortas. **C.** Acta2 transcript expression in CASC15-deficient and control mSMC. Data expressed as mean ± SEM. Student t-test.

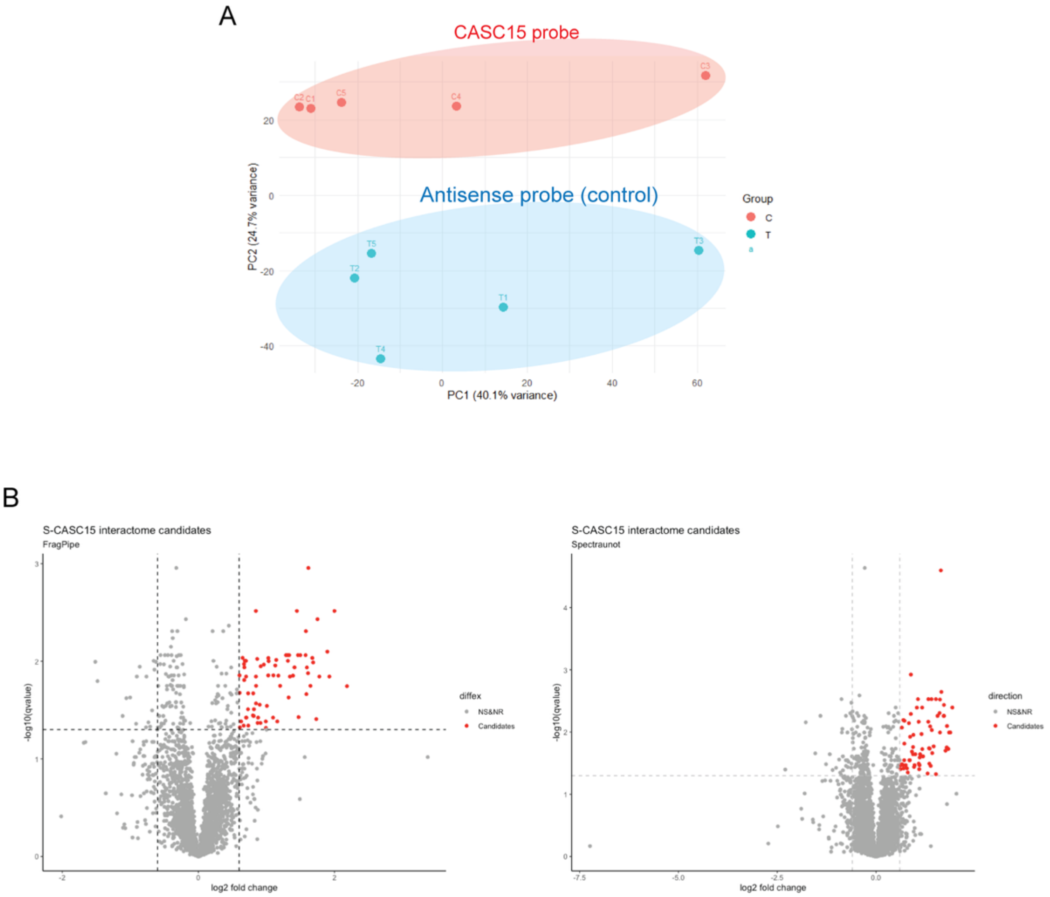


**Supplementary Figure 17: RNA pull-down paired with mass spectrometry identifies proteins interacting with CASC15. A.** PCA plot showing condition-specific clusters. **B.** Volcano plots of proteins sequenced by mass spectrometry in samples incubated with biotinylated-CASC15 probe vs biotinylated-antisense probe (control). Candidates (red dots) were selected based on q-value and fold change (q-value =0.05, $Log_2$ fold change ≥ 0.58).

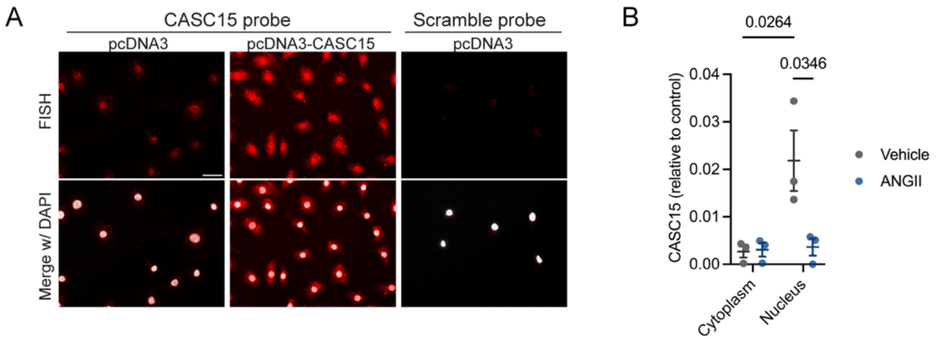


**Supplementary Figure 18: RNA-FISH reveals predominantly nuclear and perinuclear localization of CASC15 in mouse smooth muscle cells. A**. Representative CASC15 RNA-FISH (red) images in mSMCs transfected with pcDNA3 or pcDNA-CASC15. CASC15 transcripts were detected using Qiagen LNA probes (red). Nuclei were counterstained with DAPI (white). Scale bar = 50µm. **B.** CASC15 expression in mSMC cytoplasmic vs nuclear RNA extracts. SMC were treated with Angiotensin-II or vehicle for 24h.

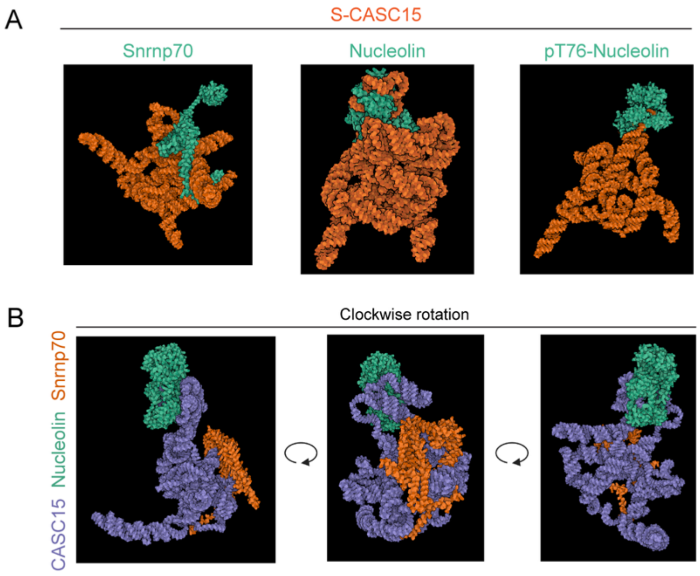


**Supplementary Figure 19: Predicted interaction between S-CASC15, Nucleolin, and SNRNP70 using Alphafold3. A.** Interaction between S-CASC15 (orange) and Snrnp70, Nucleolin, or phospho-Nucleolin (pT76-Nucleolin) (teal). **B.** Predicted formation of a CASC15 (purple), Nucleolin (teal), and Snrnp70 (orange) ternary complex.
